# Estimating total mediation effects of high-dimensional omics mediators in case-control studies

**DOI:** 10.1101/2025.01.28.635396

**Authors:** Zhiyu Kang, Li Chen, Peng Wei, Zhichao Xu, Chunlin Li, Tianzhong Yang

## Abstract

Mediation analysis helps uncover how exposures impact outcomes through intermediate variables. Traditional mean-based total mediation effect measures may suffer from the cancellation of opposite component-wise effects, and existing methods often lack the power to capture weak effects in high-dimensional mediators. Additionally, most high-dimensional mediation analysis methods have focused on continuous outcomes, with limited attention to binary outcomes, particularly in case-control studies. To fill this gap, we propose an *R*^2^ total mediation effect measure within the liability framework that offers a clear and intuitive causal interpretation, provides additional insights beyond the mean-based measures, and is invariant to disease prevalence. We develop a cross-fitted, modified Haseman-Elston regression-based estimation procedure tailored for mediation analysis in case-control studies, which can also be applied to cohort studies. Our estimator remains consistent in the presence of non-mediators and weak effects, as demonstrated in extensive simulations. Theoretical justification for consistency is provided under mild conditions and without requiring exact mediator selection. In a case-control substudy of the Women’s Health Initiative involving 2150 individuals, we found that many metabolites were mediators with weak effects in the path from BMI to coronary heart disease, and we estimated that 89% (95% CI: 57%–100%) of the BMI-explained variation in underlying CHD liability is mediated by the measured metabolomics. The proposed estimation procedure is implemented in the R package “r2MedCausal”, available on GitHub.

## 1. Introduction

Mediation analysis is a powerful tool for exploring how an exposure affects an outcome through one or more intermediate variables, known as mediators. With recent advances in high-throughput technologies, high-dimensional mediation analysis of biological data has emerged as an area of scientific interest. In this paper, we focus on estimating the total mediation effect, which provides critical insights for downstream analyses. Specifically, our method is motivated by the Women’s Health Initiative (WHI) nested case-control substudy, where we are interested in examining how obesity (quantitatively measured by the body mass index, BMI) influences the risk of incident coronary heart disease (CHD) through metabolomic profiles in postmenopausal women. Obesity (often defined as BMI ≥ 30*kg/m*^2^) is highly prevalent among adults in the US and a well-known risk factor for CHD (Eckel and Krauss, 1998). Higher BMI has been found to be associated with CHD by large epidemiological studies (Flint et al., 2010; Lyall et al., 2017; Global Burden of Metabolic Risk Factors for Chronic Diseases Collaboration et al., 2014) and causal analyses (Lyall et al., 2017; Silva, Fatumo and Nitsch, 2024). A recent randomized, double-blind, placebo-controlled clinical trial further corroborated this causal link (Packer et al., 2025), showing that a GLP-1 drug significantly reduced the risk of cardiovascular mortality and heart failure progression. With more and more weight-loss intervention strategies, understanding how BMI influences CHD risk through molecular phenotypes such as metabolomics can offer valuable insights.

Despite its importance, the mediation analysis of the WHI dataset poses a two-fold challenge. First, the existing methods fail to adequately capture the overall mediation effects, owing to the presence of many weak mediators, as suggested by our preliminary analysis in Section 6.1. A plausible reason is that many high-dimensional approaches rely on sparsity assumptions, which are not suitable for handling weak mediators with small and non-sparse effect sizes (Zhang et al., 2016; Gao et al., 2019). While some methods overcome this issue by considering transformed mediators, thus relaxing sparsity, they sacrifice interpretability (Huang and Pan, 2015). This highlights the need for a more robust, interpretable approach tailored to high-dimensional mediation analysis with weak mediators.

Second, existing mediation effect measures remain subject to ongoing improvement, especially for binary outcomes and the case-control design, despite their widespread relevance in the area (Clark-Boucher et al., 2023; Rijnhart et al., 2021). Indeed, quantifying the total mediation effect is nontrivial for high-dimensional settings. Most of the existing measures are variants of the mean-based product-of-coefficient (POC) measures, which are prone to cancellation in the presence of bidirectional mediation effects, a phenomenon that frequently occurs with omics mediators (Song et al., 2019; Yang et al., 2021). To address this issue, Yang et al. (2021) extended the variance-based *R*^2^ measure originally proposed by Fairchild et al. (2009) to accommodate multiple and high-dimensional mediators under a mixed model framework. This method captures the non-zero total mediation effect by quantifying the variance in the continuous outcome that is attributable to both the exposure and the mediators, attempting to address bidirectional effects issues in high-dimensional mediator settings. Subsequently, this measure was extended to binary outcomes using a likelihood-based McFadden’s *R*^2^ for its stability and relative independence from disease prevalence (Chi et al., 2026). However, the McFadden’s *R*^2^ defined for binary outcomes appears to be less sensitive to disease prevalence but still varies with it. Additionally, its interpretability is limited, as it is more closely linked to measuring goodness-of-fit than to explaining variance. In another line of work, Song et al. (2019) proposed a *L*_2_ norm measure of the mediation effect while simply treating binary outcomes as continuous. However, both the use of the *L*_2_ norm and the treatment of binary outcomes as continuous lack theoretical support. Importantly, all of these measures lack rigorous causal interpretations. Thus, it underscores the need for developing a mediation measure that enables a unified, robust approach to mediation analysis for binary disease outcomes, across different disease prevalence and study designs.

In response to these challenges, we propose a novel *R*^2^ mediation effect measure that provides a causal interpretation for binary outcomes. The novel *R*^2^ mediation effect measure is defined on a latent variable scale, i.e., liability, which is invariant to disease prevalence as opposed to POC measures for binary outcomes, and maintains its connection with the classic *R*^2^ defined for continuous outcomes. Thus, it offers a unified measure and a robust approach for mediation analysis for diseases with different prevalences. Unlike the traditional logistic model, the liability model requires ascertainment bias correction in a case-control study. To this end, we propose a two-stage estimation procedure to account for ascertainment bias in case-control studies and to address non-mediators and weak effects. We modify a heritability estimation method, phenotype-correlation genotype-correlation (PCGC) regression (Golan, Lander and Rosset, 2014), a generalization of Haseman-Elston regression, to estimate consistent variance components under case-control studies. It is worth noting that, although our procedure is designed for case-control studies, it is also applicable to cohort studies, thus unifying the data analysis across study designs. Extensive simulation studies showcased the robust performance of our proposed method under varying disease prevalence, study design, weak effects, and model misspecification, consistently outperforming existing high-dimensional mediation methods. Furthermore, under mild conditions and without requiring exact mediator selection, we establish the consistency of the estimators. Applied to the motivating WHI case-control dataset, our method effectively captures a good amount of mediation effect of metabolites from BMI to CHD.

The rest of the paper is organized as follows. Section 2 reviews the existing mediation effect measures and highlights their limitations. Section 3 introduces our proposed 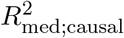 mediation measure and discusses its causal interpretation. Section 4 details the estimation procedure and addresses implementation considerations specific to case-control designs. Section 5 evaluates the performance of our method through extensive simulation studies, comparing it with existing approaches. Section 6 presents the application to WHI. Section 7 summarizes the contributions and limitations of our work and outlines future research directions. The theoretical proofs are provided in Section 1 of the Supplementary Material (Kang et al., 2025). Finally, an R package, “r2MedCausal”, has been developed to implement the proposed estimation procedure.

## 2. Preliminaries

This section provides an overview of existing approaches and their limitations, which serve as the basis for the proposed approach.

A foundational framework in mediation analysis is established by Baron and Kenny (1986), who proposed the following structural equations for a continuous outcome, describing the relationships among an exposure *X*, a *p*-dimensional vector of mediators ***M***, and an outcome *Y* :

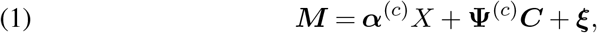

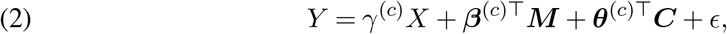

where ***C*** is a *q ×* 1 vector of covariates. The superscript ^(*c*)^ decorates coefficients corresponding to the continuous outcome setting. The coefficient 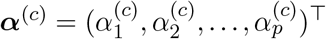 captures the exposure-mediator relationship after adjusting for the covariates. Similarly, 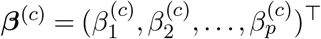 captures the mediator-outcome relationship after adjusting for both exposure and covariates. The parameter *γ*^(*c*)^ is the direct effect of the exposure variable, and **Ψ**^(*c*)^ and ***θ***^(*c*)^ are the coefficients of covariates. The variables *ϵ*, ***ξ*** = (*ξ*_1_, *ξ*_2_, …, *ξ*_*p*_)^⊤^ are error terms. In model (1)–(2), the total effect of the exposure *X* on the continuous outcome *Y* is decomposed into a direct effect and a mediation effect. The total effect ***α***^(*c*)⊤^***β***^(*c*)^ + *γ*^(*c*)^ is typically estimated by regressing *Y* on *X*, after adjusting for covariates. The mediation effect can be estimated using either the difference-in-coefficients method, which subtracts the direct effect *γ*^(*c*)^ from the total effect, or the POC approach, which computes the inner product ***α***^(*c*)⊤^***β***^(*c*)^.

Most early development of mediation analysis was not formulated in a rigorous causal framework (Pearl, 2001; VanderWeele and Vansteelandt, 2010). In causal mediation analysis (with a continuous outcome), the natural indirect effect (NIE) and the natural direct effect (NDE) for an exposure contrast between *x* and *x*^*′*^ are defined as:

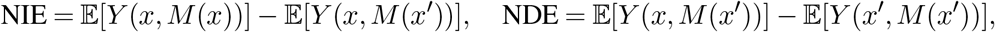

where *Y* (*x, M* (*x*)) is the potential outcome when both exposure and mediator take their natural values under *X* = *x, Y* (*x, M* (*x*^*′*^)) is the outcome when the exposure is set to *x*, but the mediator is fixed at the level it would take under *X* = *x*^*′*^, and *Y* (*x*^*′*^, *M* (*x*^*′*^)) represents the outcome when both exposure and mediator take their natural values under *X* = *x*^*′*^. To identify NIE and NDE, the following assumptions are often imposed (VanderWeele, 2016).

### Assumption 1

(No Unmeasured Confounding).

**A1** No unmeasured confounding of the exposure-mediator effect.

**A2** No unmeasured confounding of the mediator-outcome effect.

**A3** No unmeasured confounding of the exposure-outcome effect.

**A4** No mediator-outcome confounder that is itself affected by the exposure.

If Assumption 1 holds in model (1)–(2), then we have NIE = ***α***^(*c*)⊤^***β***^(*c*)^(*x* − *x*^*′*^), which provides POC with a rigorous causal interpretation.

### 2.1. The issues of POC and its alternatives

Partly due to its causal interpretation, the POC measure and its variants, such as the proportion-mediated (PM) measure ***α***^⊤^***β****/*(*γ* + ***α***^⊤^***β***) (MacKinnon et al., 2007), have been popular, including in high-dimensional mediation analysis of biological data (Zhang et al., 2016; Gao et al., 2019). However, only reporting POC-based measures can be misleading in high-dimensional settings where numerous bidirectional component-wise mediation effects (i.e., *α*_*i*_*β*_*i*_ have different signs) are present (Song et al., 2019), potentially causing cancellation in the POC and PM measures. Therefore, a small (or near-zero) POC may not imply the absence of mediation, but rather that positive and negative indirect pathways offset in aggregate.

Recognizing these drawbacks, Song et al. (2019) proposes an alternative *L*_2_ norm for summarizing the overall mediation effects: 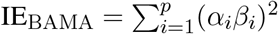. While this modification no longer suffers from cancellation of bidirectional component-wise mediation effects, it lacks rigorous theoretical support for the causal interpretation. An alternative mediation effect measure is the *R*^2^ measure, which was first proposed in a single-mediator model for continuous outcomes by Fairchild et al. (2009), and later extended by Yang et al. (2021) to high-dimensional mediation analysis:

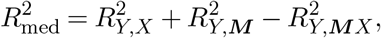

where 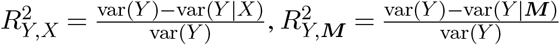, and 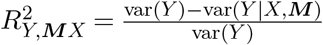. The *R*^2^ measure provides a robust, sensible variance-based measure for continuous outcomes. However, it does not have a clear causal interpretation.

### 2.2. The lack of a unified, robust measure for binary outcomes

The mediation framework is extended for binary outcomes. This is often done by replacing (2) with a logistic regression (MacKinnon et al., 2007). In this situation, the difference-in-coefficients method may encounter the noncollapsibility problem (Greenland, Pearl and Robins, 1999), making the POC approach preferable (VanderWeele, 2016). When Assumption 1 is satisfied and the outcome is rare, logistic regression can be used to estimate the causal mediation effect on an odds ratio scale as: OR^NIE^ ≈ exp(***α***^⊤^***β***) (VanderWeele and Vansteelandt, 2010). However, when the outcome is not rare, odds ratios suffer from noncollapsibility, meaning that exp(***α***^⊤^***β***) does not provide an interpretable mediation effect estimate, as it depends on covariates adjustment. In such cases, log-binomial models are common to estimate the causal mediation effect on a risk ratio (RR) scale, given by RR^NIE^ ≈ exp(***α***^⊤^***β***) (VanderWeele, 2016). Consequently, the preferred mediation effect scale varies by outcome prevalence, with odds ratios for rare outcomes and risk ratios for common ones. Moreover, the foregoing approaches are fundamentally based on the POC calculation, thus suffering from the same cancellation problem as described for continuous outcomes earlier.

Motivated by our analysis of WHI data, we aim to develop a new measure for mediation analysis of binary outcomes, which enjoys the following merits: (i) it has a causal interpretation; (ii) it overcomes the cancellation issue; and (iii) it offers a unified treatment for binary outcomes with different prevalence and study designs.

## 3. Modeling

For binary outcomes, the liability model provides a unified approach for analyzing both rare and common diseases. In addition, the variance-based mediation effect has a consistent interpretation for both continuous outcomes and binary outcomes under the liability model. It has been scarcely used in low-dimensional settings (MacKinnon et al., 2007; Nguyen et al., 2016) and recently adapted to high-dimensional settings (Gaynor, Schwartz and Lin, 2019; Sohn, Lu and Li, 2022). Thus, we utilize the liability-based mediation model, which is defined by the following structural equations:

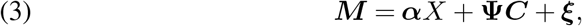

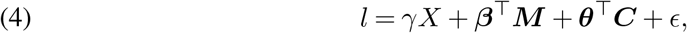

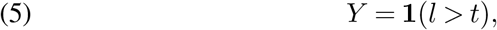

where *l* is the underlying latent variable, *Y* is the observed binary outcome. The function **1**(·) is an indicator function, with threshold *t* determining *K*, the prevalence of the trait in the population. The coefficients ***α, β***, *γ*, **Ψ**, and ***θ*** retain the same interpretations as in the continuous outcome model, now defined in relation to the latent liability. The error terms *ϵ* and ***ξ*** are assumed to be normally distributed.

In what follows, we perform analyses conditioned on ***C*** and treat ***C*** as fixed. Additionally, without loss of generality, *X* is assumed to be centered.

### 3.1. Modeling non- and weak mediators

In real-world data, the identities of true mediators are usually unknown, necessitating the inclusion of non-mediators in our mediation model. Potential mediators ***M*** are partitioned into true mediators ***M***_𝒯_, and three types of non-mediators (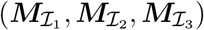), as illustrated in Figure 1:

- ***M***_𝒯_ = {*M*_*j*_|*α*_*j*_ ≠ 0 and *β*_*j*_ ≠ 0 for *j* ∈ 𝒯 };
- 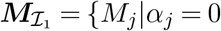 and *β*_*j*_ ≠ 0 for *j* ∈ ℐ_1_};
- 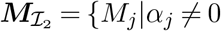 and *β*_*j*_ = 0 for *j* ∈ ℐ_2_};
- 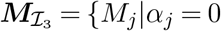 and *β*_*j*_ = 0 for *j* ∈ ℐ_3_}.

**Fig 1.**
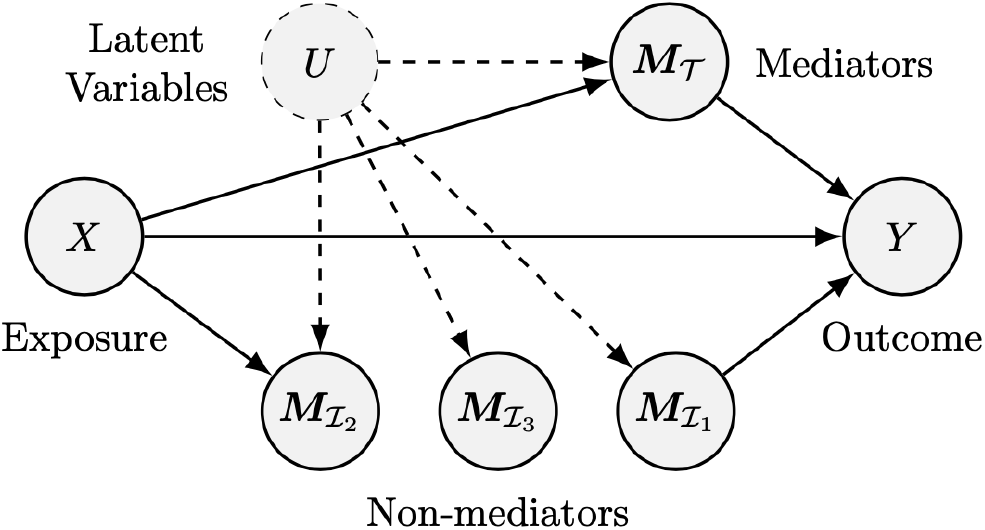
Graph representation of a mediation model.

We denote the proportions of ***M***_𝒯_, 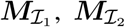, and 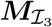 as *π*_11_, *π*_01_, *π*_10_ and *π*_00_ respectively. The overall proportions of non-zero elements in ***α*** and ***β*** are then given by *π*_*α*_ = *π*_11_ + *π*_10_ and *π*_*β*_ = *π*_11_ + *π*_01_.

Existing approaches often assume the true mediators are sparse (*π*_11_ ≈ 0) and try to identify mediators through either high-dimensional sparse linear regression (Zhang et al., 2016; Gao et al., 2019) or large-scale hypothesis testing (Huang and Pan, 2015). The success of selection roughly requires the signal strength |*α*_*k*_|∧ | *β*_*k*_| to be larger than 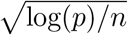. However, such approaches may be plagued by the presence of dense *weak mediators* such that 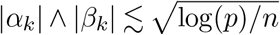.

Moving beyond the sparse regime, we model the dense mediation effects based on the random-effects hypothesis (Dobriban and Wager, 2018). Heuristically, the results obtained under this random-effects hypothesis can be regarded as average-case analyses over dense parameters, offering a qualitatively different perspective for high-dimensional mediation analysis than the popular sparse regression approaches. Specifically, we consider a mixed-effect model under the working assumptions that

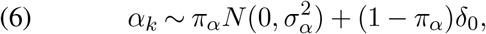

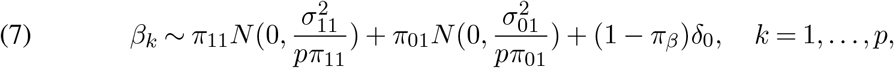

where *δ*_0_ denotes a point mass at zero. The purpose of introducing the normalizing factor *p*^−1^ in (7) is to let the per-mediator contribution vanish and thus the aggregated mediation variation remains finite (Dobriban and Wager, 2018). Consequently, it becomes impossible to accurately select mediators in this scenario, as we have 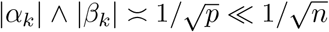; nevertheless, it remains feasible to estimate the aggregated mediation variation.

### 3.2. A novel R^2^ measure with causal interpretation

Aiming to achieve the robustness of rare disease assumption and ease of interpretation, we define a new *R*^2^ measure for high-dimensional mediators under the liability scale for binary outcomes. Following Yang et al. (2021), under the structural equation model specified by (3)–(5), the *R*^2^ measure would be defined as:

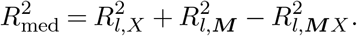

However, this definition of 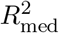 does not inherently take into account the directional relationship between *X* and ***M***. Specifically, exchanging *X* and ***M*** would not change 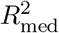, despite mediation analysis explicitly assuming directed causal pathways from *X* to *Y* and from *X* to ***M*** to *Y*. To resolve this issue and explicitly incorporate the causal direction, we introduce the do-operators in our *R*^2^ measure within Pearl’s Structural Causal Framework (Pearl, 2010). We formulate the new *R*^2^ measure for binary outcomes as:

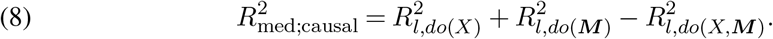

Based on (3)–(5) and working model assumptions (6)–(7), we define the *causal coefficient of determination* as

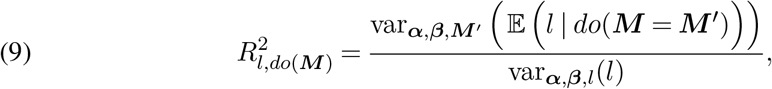

where ***M*** ^*′*^ is identically distributed as ***M*** but independent of all other variables in the model and the do-operator is a stochastic intervention of ***M***. The stochastic intervention *do*(***M*** = ***M*** ^*′*^) induces the so-called truncated factorization (Pearl, 2010), which removes all the incoming edges of ***M*** in Figure 1 and replace ***M*** with a new random vector ***M*** ^*′*^. Thus, 𝔼 (*l*|*do*(***M*** = ***M*** ^*′*^)) is the (counterfactual) mean of *l* conditioned on ***M*** ^*′*^ with respect to the truncation probability, had ***M*** been replaced by ***M*** ^*′*^. In causal mediation model (3)–(5), 𝔼 (*l* | *do*(***M*** = ***M*** ^*′*^)) = ***β***^⊤^***M*** ^*′*^ + ***θ***^⊤^***C*** + 𝔼 (*γX* + *ϵ*) = ***β***^⊤^***M*** ^*′*^ + ***θ***^⊤^***C*** (because we treat ***C*** as fixed and *X* is centered). As a result, 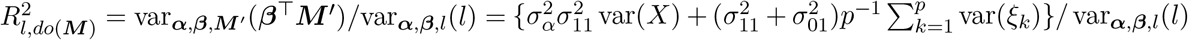. By introducing the stochastic intervention, it ensures that 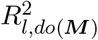 reflects the causal effect of ***M*** on *l*, in contrast to 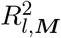, which may be confounded by *X*. Similarly, we have

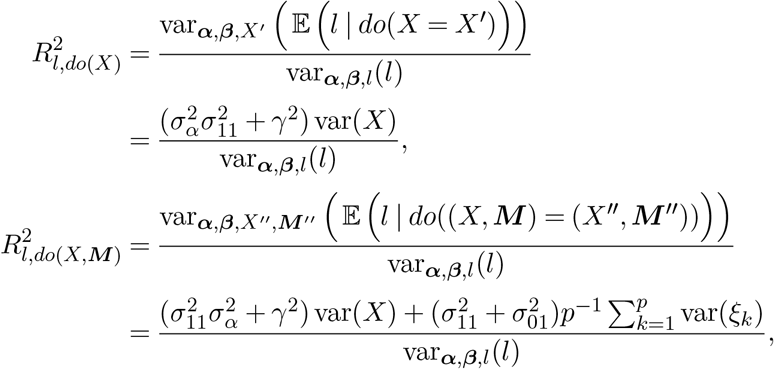

where *X*^*′*^ and (*X*^*′′*^, ***M*** ^*′′*^) are independent copies of *X* and (*X*, ***M***), respectively, drawn from the same distribution but independent of other variables in the model. Thus, we have

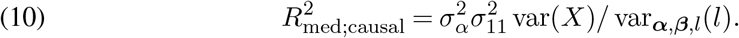

The proposed 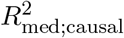 can be interpreted as the proportion of variation in the underlying liability for the outcome that is *causally* explained by the exposure through the mediators.

As discussed in MacKinnon et al. (2007), the PM measure in traditional mediation analysis is given by ***α***^⊤^***β****/*(*γ* + ***α***^⊤^***β***). In the causal mediation analysis framework, assuming a rare binary outcome, the PM measure can be expressed as OR^NDE^(OR^NIE^ − 1)*/*(OR^NDE^OR^NIE^− 1), where OR^NDE^ = exp (*γ*) and OR^NIE^ = exp(***α***^⊤^***β***) denote the NDE and NIE on the odds ratio scale (VanderWeele, 2016). However, neither measure is guaranteed to range from 0 to 1. Additionally, both rely on the sum of component-wise effects, suffering from the cancellation problem when effects are in different directions. As an alternative to the PM measure, we propose a new relative 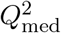 measure, which is bounded between 0 and 1:

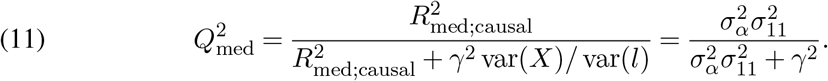

This quantity is the proportion of variation of liability causally explained by *X* indirectly through the mediators over the sum of the proportion of the variation explained indirectly and directly by *X*. It can be interpreted as the proportion of the exposure explained variance on the liability scale of the outcome that is accounted for by the mediators.

#### Remark 1

(Relationship with causal entropy). The proposed causal coefficient of determination is connected to the existing concept of causal entropy (Simoes, Dastani and van Ommen, 2024). It can be shown that for a single mediator,

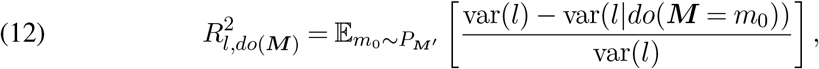

where *do*(***M***) = *m*_0_ implies that we intervene ***M*** and set it to be *m*_0_, ignoring the dependencies between *M* and its parent variables. This notation explicitly accounts for causality (direction) in our formula. It can be shown that (9) and (12) are equivalent for a single mediator. The connection between our definition and causal entropy can be seen directly through (12).

## 4. Estimation procedure

To estimate the proposed *R*^2^ measure, we adapted the cross-fitting framework as described by Xu et al. (2024) to improve the robustness of mediation effect estimation. Cross-fitting mitigates bias introduced by the winner’s curse during the variable screening step (Göring, Terwilliger and Blangero, 2001). The approach involves a two-stage procedure. First, the sample is randomly split into two subsamples 𝒟^(1)^ and 𝒟^(2)^. In Step 1, mediation selection and variance estimation of ***α*** are performed on subsample 𝒟^(1)^ to get the filtered mediator index set 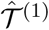 and variance estimate 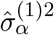. In Step 2, the estimation of other parameters is conducted on subsample 𝒟^(2)^ based on the selected mediators 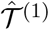. The roles of the two subsamples are then reversed, allowing the whole sample to be used in the estimation process. Finally, the *R*^2^ estimate is given by the mean of cross-fitted *R*^2^ estimates 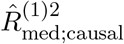 and 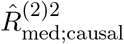. The procedure is visualized in Section 4 of the Supplementary Material (Kang et al., 2025), and the mediator selection and parameter estimation process are detailed in Subsections 4.1-4.3.

### 4.1. Estimation of 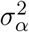 and mediator selection

The estimation of ***α*** in our mediation analysis framework has been extensively studied as a secondary outcome analysis problem in case-control studies (Schifano, 2019). To estimate 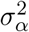, we used the inverse probability weighting (IPW) method introduced by Horvitz and Thompson (1952). In this approach, each individual’s contribution to the estimating equations is weighted by the reciprocal of their selection probability, effectively constructing a pseudo-sample that is more representative of the target population than the original case-control sample.

Let *S*_*i*_ denote the inclusion indicator for the *i*-th individual, and define the inverse weight as *w*_*i*_ = 1*/P* (*S*_*i*_ = 1|*Y*_*i*_, ***M***_*i*_, *X*_*i*_, ***C***_*i*_). The IPW estimating equation for estimating each element of ***α*** is given by (Sofer et al., 2017):

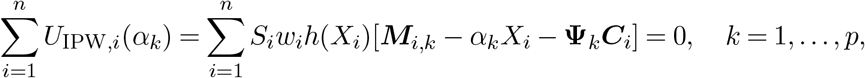

where **Ψ**_*k*_ is the *k*-th row of coefficient **Ψ**, *h*(*X*_*i*_) is a user-specified function ensuring that 𝔼 (*∂U*_IPW,*i*_*/∂α*_*k*_) is invertible. Under the full-ascertainment assumption (Stene, 1977), *w*_*i*_ = 1 for case and 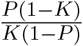 for control, where *K* represents the population-level trait prevalence, and *P* denotes the case-control ratio in the study, both assumed to be known. Setting *h*(*X*_*i*_) = *X*_*i*_ simplifies the estimating equation into a weighted least squares regression problem.

To identify non-mediators and noise variables, Wald tests are conducted to detect 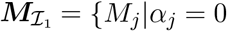 and *β*_*j*_ = 0 for *j* ∈ ℐ_1_} and 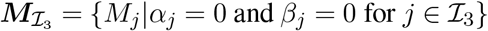. Wald test p-values are adjusted using a multiple testing correction procedure to control the false discovery rate (FDR) at a pre-specified threshold. The resulting filtered set of mediators, denoted by 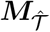, is subsequently used in cross-fitted variance component estimation. Given that 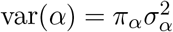, the variance estimator is obtained as 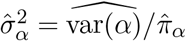, where var(*α*) is empirically estimated from the IPW results, and *π*_*α*_ is estimated based on the ratio of the number of selected mediators and the total number of potential mediators.

### 4.2. Estimation of γ

The IPW approach can also be applied to estimate the parameter *γ*. By selecting an appropriate estimating equation, the IPW problem becomes a weighted probit regression problem with the same weight as described in Section 4.1. However, directly regressing the outcome *Y* on the exposure variable *X* and covariates ***C*** may lead to biased estimates, as it does not account for ***M***. To mitigate this bias, we include the top principal components (PCs) of the mediator matrix across individuals, denoted as ***W***, as additional covariates in the model. By incorporating these components, we can effectively control for the influence of ***M*** that may introduce bias in the estimation of *γ* (Lemma 3, Section 1 of the Supplementary Material (Kang et al., 2025)).

### 4.3. Estimation of 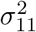

For the estimation of 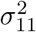, we propose to use PCGC regression (Golan, Lander and Rosset, 2014). PCGC regression is a method of moments that estimates variance components under a liability model while accounting for the oversampling in case-control designs. As a generalization of the Haseman–Elston regression method (Haseman and Elston, 1972), PCGC regression models the expected correlation of the observed phenotype as a function of variance components and the correlation of its corresponding design matrix, which, in our context, corresponds to the mediator matrix across individuals. Unlike the restricted maximum likelihood (REML) method (Lee et al., 2011), which tends to underestimate variance components in case-control studies due to ascertainment bias, PCGC explicitly accounts for this bias by modeling the inclusion probability of individuals in the study. However, this approach implicitly assumes weak correlation among rows of the design matrix, an assumption that is likely violated by mediators due to their relationship with the exposure (Equation (3)). To account for this, we modify the PCGC regression framework and estimate 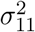 following the cross-fitting procedure as described in Section 4:

Step 1 (First subsample): Identify the mediators with *α*≠ 0, i.e., ***M***_𝒯_ and 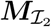 following Section 4.1 and denote the selected mediator set as 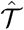.

Step 2 (Second subsample): Estimate standardized residual 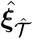 by regressing scaled ***M*** on *X*, ***C, W*** using the IPW approach. Apply PCGC regression with 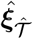 as the standardized design matrix and include *X*, ***C*** and ***W*** as covariates to estimate 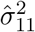.

After adjusting for covariates that drive the correlation between mediators, the resulting residuals 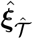 in Step 2 would satisfy the weak correlation assumption in PCGC regression. We note that adjusting for ***W*** in the analysis not only provides us a consistent estimate of direct effect in high-dimensional settings (Section 4.2), but also potentially controls for unknown confounding in the same spirit as Xu et al. (2024). Additionally, ascertainment bias is appropriately addressed in both steps through the IPW approach for residual estimation and PCGC regression for variance estimation.

### 4.4. Theoretical results on consistency

In this section, we present the theoretical properties of the proposed method. For theoretical analysis, we assume that while the mediators may be correlated, there are no directed causal relationships among them. Instead, their correlation is driven by the exposure *X*, covariates ***C***, and unobserved/unknown latent factors ***U*** (Figure 1). This assumption is formalized as follows.

#### Assumption 2

(Conditional Parallel Mediators).

There exist random variables ***U*** that have a global influence on ***M***, where *M*_*k*_ ╨ *M*_*j*_|*X*, ***U***, ***C*** for any *k, j* ∈ {1, …, *p*}.

Assumption 2 is commonly used in methods and applications to high-dimensional omics mediation analysis (Huang and Pan, 2015; Perera et al., 2022; Chi et al., 2026), as usually the omics data were measured at a single time point rather than sequentially, exemplified by our motivating dataset, WHI. An example for such unobserved/unknown latent variables ***U*** is batch effect (e.g., lab conditions, technicians, etc.), which frequently occurs in biological data and typically happens after sample collection (i.e., ***U*** ╨ *S*). In our proposed estimation procedure, the random variable ***U*** is captured by PC analysis (Section 4.2). Moreover, our estimation method remains robust when Assumption 2 is violated, as empirically shown in Supplementary Material Section 2.8 (Kang et al., 2025).

Given Assumptions 1 and 2, Theorem 4.1 establishes the consistency of our proposed total mediation effect estimator 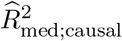 and relative measurement estimator 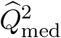 under mild regularity conditions.

#### Theorem 4.1.

*Under Assumptions 1-2 and Assumptions 3-10 in Section 1 of the Supplementary Material (Kang et al., 2025), and assume there exists c*_0_ *>* 0 *such that* 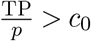, *then for any ϵ >* 0, *there exists δ >* 0, *such that if* 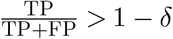 *for p, n large enough, we have*

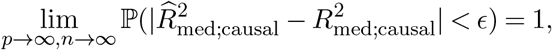

*and*

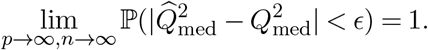

Let 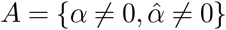 and 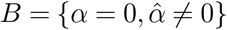. Herein, TP and FP correspond to |*A*| and |*B*| in the initial filtering step of *α* (Section 4.1). A key assumption underlying this result is that the proportion of true mediators and non-mediators type 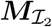 among all potential mediators remains above a positive threshold as *p →* ∞. This assumption ensures a sufficiently large number of mediators contribute to the total mediation effect. Another important assumption is the control of FDR when identifying non-mediators type 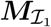 and noise mediators 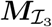. We assume that the FDR for selecting variables impacted by exposure is stringent enough to minimize the inclusion of non-mediators type 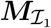 and noise mediators 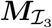. In practice, our simulations suggest that a reasonable choice, such as FDR = 0.01 using Benjamini-Hochberg (BH) procedure, yields robust performance across various sample sizes and mediator sparsity levels (Detailed empirical evaluation in Supplementary Section 2.7). Notably, the proposed estimator is shown to be consistent without requiring mediator selection consistency but only sufficiently low FDR in testing *p* marginal association (i.e., *α* = 0). This is especially desirable in high-dimensional situations, as exact selection is rarely achieved.

## 5. Simulation studies

### 5.1. Simulation settings

We conducted simulation studies to evaluate the consistency of the proposed estimators of 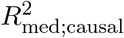 and 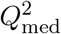. Four scenarios were considered, each varying the composition of mediator types while fixing the number of mediators *p* = 2000:

- Scenario 1 (S1): 180 ***M***_𝒯_, 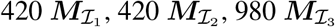;
- Scenario 2 (S2): 360 ***M***_𝒯_, 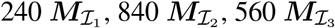;
- Scenario 3 (S3): 360 ***M***_𝒯_, 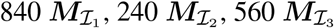;
- Scenario 4 (S4): 720 ***M***_𝒯_, 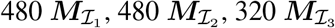.

The binary outcome was generated under the liability threshold model, with population disease prevalence *K* = 0.05 and study disease prevalence *P* = 0.5 to mimic a case-control design with a 1:1 case-control ratio. Sample size *n* varies from 500 to 2000 with an increment of 500. We set a single exposure variable *X* ∼ *N* (0, 1) and its direct effect parameter *γ* = 0.5. We specified 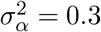, the cumulative variance of the true mediators ***M***_𝒯_ as 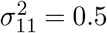, and that of the non-mediators 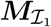 as 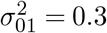. The coefficients ***α*** and ***β*** were generated from the working model (6). To ensure Assumptions 1 and 2 hold, we generated *ξ*_*k*_ = *α*_0_*U* + *ν*_*k*_ for *k* = 1, …, *p*, where *U* ∼ *N* (0, 1), *ν*_*k*_ ∼ *N* (0, 1) and *α*_0_ ∼ *N* (0, 0.3), with *U, ν*_*k*_ and *α*_0_ mutually independent. The vector of ***ξ*** = (*ξ*_1_, …, *ξ*_*p*_)^⊤^ was then used to generate mediators as ***M*** = ***α****X* + ***ξ***. The error term for the outcome model was generated from 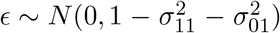. Additionally, the simulation settings under the global null, i.e., no mediators and no mediation effects, are described in Section 2.1 of the Supplementary Material (Kang et al., 2025). For each scenario, we ran 100 replications across all sample sizes.

Under S1, we further compared our methods to competing methods: HIMA (Zhang et al., 2016), HDMA (Gao et al., 2019) and BAMA (Song et al., 2019) in terms of total mediation effect estimate and PM estimate. We varied the prevalence of disease *K* from 0.5 (cohort study) to 0.05 (case-control study) and the sample size from 500 to 2000. For the competing methods, we calculated mean-based mediation effects as recommended in their paper. For methods HIMA and HDMA, their IE measure is the inner product of 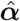 and 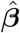. Their proportion mediated, following the definition in traditional analysis, is 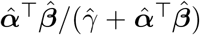. While for BAMA, the mediation effect measure is given by 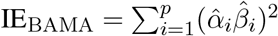. The results are summarized in the following section. Furthermore, we calculated the corresponding 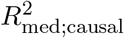 and 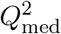 based on their component-wise estimates 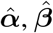 and 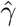. Relative bias was also reported for fair comparisons across different measures, defined as Bias*/*True Value.

### 5.2. Simulation results

Figure 2 and Table 1 summarize the estimation performance of two proposed measures in terms of bias, standard deviation, mean absolute deviation and mean square error. Overall, the proposed measures had minimal bias and error observed across different sample sizes and number/types of non-mediators. As expected, the performance metrics improved as the sample size increased. Furthermore, the violin plots (Figure 2) indicate the consistency of the proposed estimation methods as well as their robustness under various conditions. The performance of our proposed estimators under the global null is shown in Section 2.1 of the Supplementary Material (Kang et al., 2025), where all performance metrics are at the same scale as the Table 1, suggesting consistency in both null and alternative cases.

**Fig 2.**
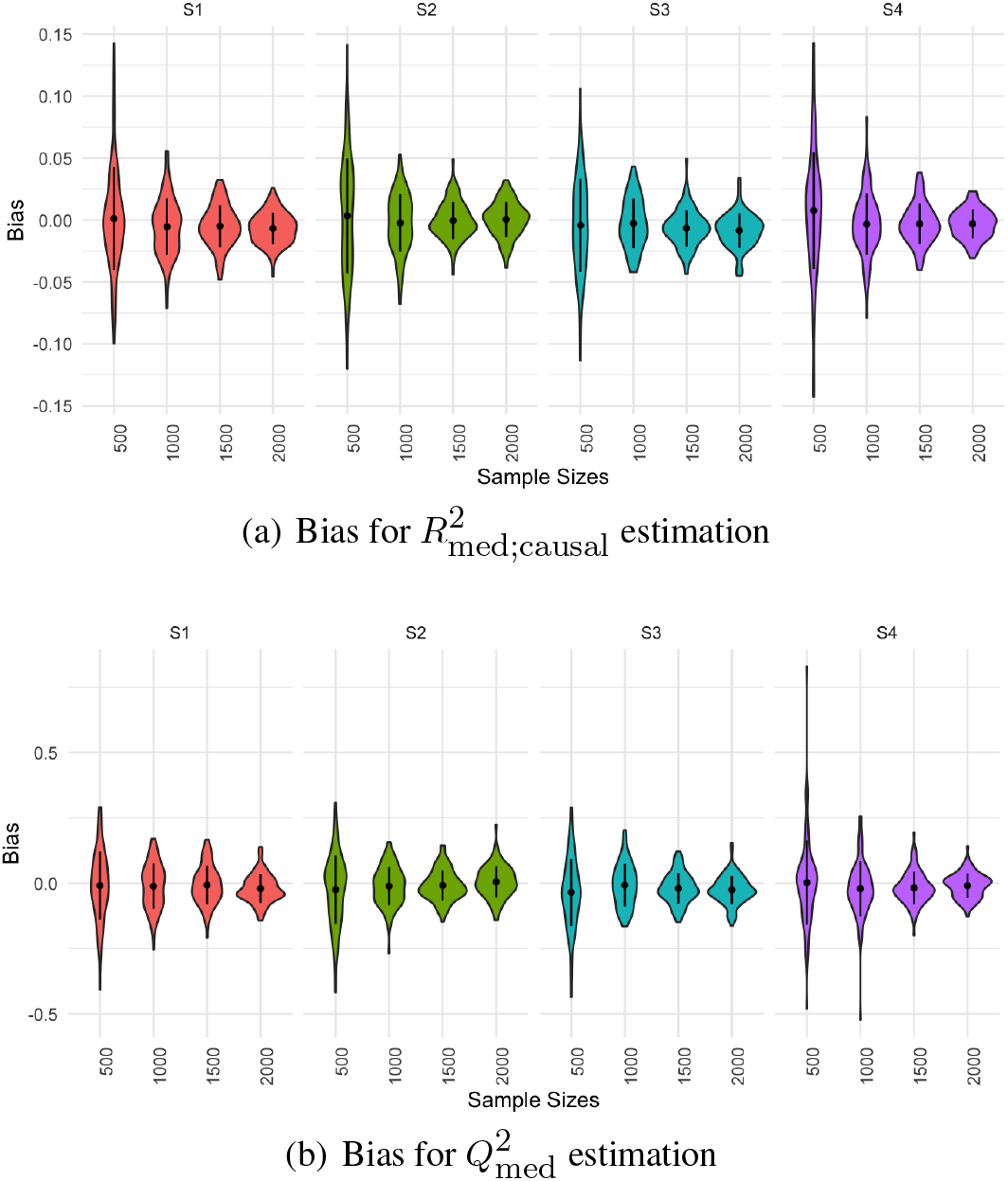
Violin plots of bias in a case-control study (K = 0.05) under varying sample sizes and proportions of non-mediators across Scenarios S1-S4. The black dots represent the average bias, while the vertical lines correspond to the range of one standard deviation centered at the mean bias.

**Table 1.** Performance of proposed estimators under different scenarios. Bias represents empirical bias, SD represents standard deviation, MAD is mean absolute deviation, and MSE is mean square error, all based on 100 repetitions.

| Scenario | $n$ | $R^2_{\text{med;causal}}$ | | | | $Q^2_{\text{med}}$ | | | |
| --- | --- | --- | --- | --- | --- | --- | --- | --- | --- |
|  |  | Bias | SD | MAD | MSE | Bias | SD | MAD | MSE |
| S1 | 500 | 0.0012 | 0.0415 | 0.0297 | 0.0017 | -0.0088 | 0.13 | 0.101 | 0.0167 |
|  | 1000 | -0.0054 | 0.0229 | 0.0183 | 0.0005 | -0.0114 | 0.0864 | 0.0714 | 0.0075 |
|  | 1500 | -0.0049 | 0.0168 | 0.0139 | 0.0003 | -0.0071 | 0.0735 | 0.0592 | 0.0054 |
|  | 2000 | -0.0067 | 0.0128 | 0.0115 | 0.0002 | -0.0209 | 0.0543 | 0.0469 | 0.0034 |
| S2 | 500 | 0.0035 | 0.0464 | 0.0387 | 0.0021 | -0.0248 | 0.132 | 0.103 | 0.0177 |
|  | 1000 | -0.0023 | 0.0232 | 0.0189 | 0.0005 | -0.011 | 0.0734 | 0.0592 | 0.0055 |
|  | 1500 | -0.0003 | 0.0150 | 0.0116 | 0.0002 | -0.0084 | 0.0583 | 0.0469 | 0.0034 |
|  | 2000 | 0.0006 | 0.014 | 0.0111 | 0.0002 | 0.0053 | 0.0603 | 0.047 | 0.0036 |
| S3 | 500 | -0.0042 | 0.0377 | 0.0302 | 0.0014 | -0.0351 | 0.128 | 0.104 | 0.0175 |
|  | 1000 | -0.0027 | 0.0201 | 0.0168 | 0.0004 | -0.0072 | 0.0813 | 0.0666 | 0.0066 |
|  | 1500 | -0.0065 | 0.0147 | 0.0125 | 0.0003 | -0.0197 | 0.0579 | 0.0492 | 0.0037 |
|  | 2000 | -0.0084 | 0.0139 | 0.0124 | 0.0003 | -0.0258 | 0.054 | 0.0469 | 0.0035 |
| S4 | 500 | 0.0077 | 0.0471 | 0.0359 | 0.0023 | 0.0025 | 0.162 | 0.108 | 0.0259 |
|  | 1000 | -0.0033 | 0.0248 | 0.0188 | 0.0006 | -0.021 | 0.105 | 0.0771 | 0.0114 |
|  | 1500 | -0.0031 | 0.0162 | 0.013 | 0.0003 | -0.018 | 0.0618 | 0.0505 | 0.0041 |
|  | 2000 | -0.0031 | 0.0118 | 0.0098 | 0.0001 | -0.0095 | 0.0466 | 0.0371 | 0.0022 |

Table 2 presents a comparison of our method with HIMA, HDMA, and BAMA in terms of bias, standard deviation, and mean square error for estimating IE and PM under case-control designs (*K* = 0.05). Our proposed method consistently outperformed competing methods across all simulation scenarios. HIMA and HDMA often identified only a small number of mediators (typically fewer than 10), which indicated their limited ability to capture mediators with small effect sizes, resulting in higher bias and less reliable estimates. Moreover, neither HIMA nor HDMA accounts for ascertainment bias when estimating the exposure–mediator associations ***α***, which likely contributes to their biased effect estimates under case-control designs. BAMA, on the other hand, explicitly assumes that mediator effects follow a normal distribution with either large or small variance, which may explain its improved performance over HIMA and HDMA in some simulation scenarios. However, the amount of bias was still higher than desired. Simulation results with *K* = 0.2 and 0.5 are presented in Section 2 of the Supplementary Material (Kang et al., 2025). As the prevalence of disease *K* increased, our method showed reduced efficiency in small sample sizes, indicated by the higher estimation variance. However, with larger sample sizes, the method’s performance became more stable, yielding comparable results across different *K* values.

**Table 2.**
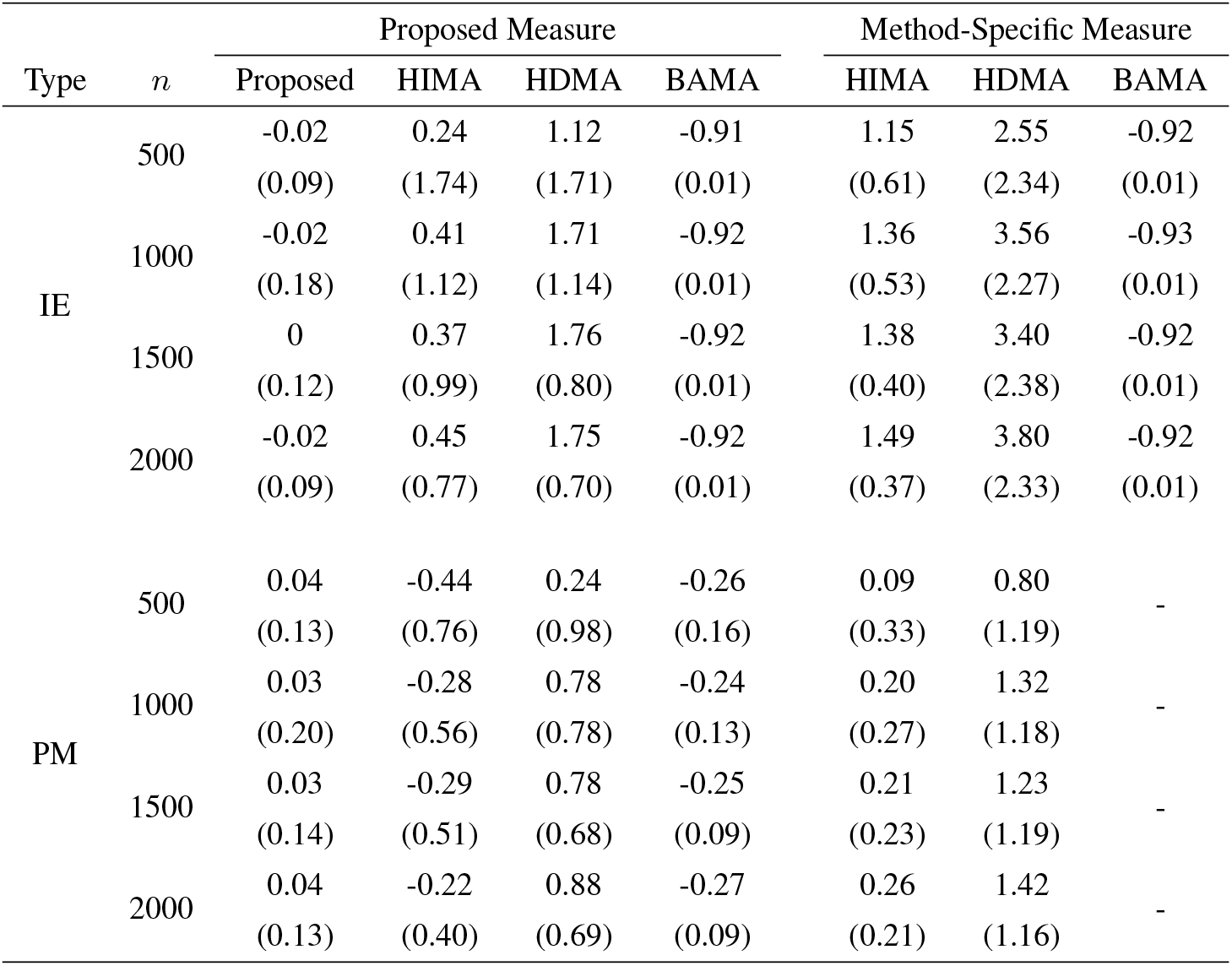
Comparison of the relative performance of IE and PM estimators in a case-control study (K = 0.05). Results are shown for both the proposed measures (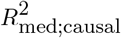 and 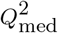) and the original method-specific measures: POC estimator for HIMA and HDMA, and 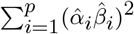 for BAMA when estimating IE; and 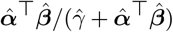 for HIMA and HDMA when estimating PM. Each cell presents the relative bias, with the standard deviation of the relative deviation shown in parentheses

Computation times in both case-control and cohort studies are summarized in Section 2.3 of the Supplementary Material (Kang et al., 2025). No significant difference in running time was observed between cohort (*K* = 0.5) and case-control (*K* = 0.05) studies across all methods. However, the running time of other methods increased noticeably with larger sample sizes. Overall, HIMA, HDMA, and BAMA exhibited approximately linear growth as *n* increased, while the proposed method has the slowest growth in running time. Particularly, BAMA, under a Bayesian sampling estimation framework, is the slowest approach. While HDMA was faster than BAMA, it remained slower than HIMA and the proposed method.

In Supplementary Sections 2.4–2.8 (Kang et al., 2025), we further assess the robustness of our method in the presence of a few strong mediators, disease prevalence misspecification, small total effects, varying FDR thresholds, and violation of the parallel mediation assumption. Overall, the proposed approach consistently demonstrated stable performance across these realistic scenarios.

## 6. Application to the Women’s Health Initiative (WHI) study

The WHI nested case-control substudy (dbGaP accession number: phs001334) examines the relationship between metabolomic profiles and incident CHD in postmenopausal women. The substudy consists of subjects from both the WHI Observational Study (WHI-OS) and the WHI Hormone Therapy Trials (WHI-HT). A total of 2,306 samples (681 case-control pairs from WHI-HT and 472 case-control pairs from WHI-OS) were analyzed for metabolomic profiling, representing 2,151 unique consented participants in the analysis. Baseline targeted metabolomic profiles were measured using liquid chromatography–mass spectrometry, yielding 371 metabolites after initial quality control. Cases were defined as postmenopausal women who experienced incident myocardial infarction (MI) or CHD death during follow-up. In WHI-HT, cases were those who developed MI or CHD death during the hormone therapy intervention period. In WHI-OS, cases were selected from previously identified incident CHD cases in the prior biomarker study (BAA 22), excluding those who experienced a stroke before MI or CHD death. Controls were matched by age, race/ethnicity, enrollment period, and hysterectomy status, excluding those with prior cardiovascular diseases.

BMI is a well-established modifiable risk factor for CHD (Eckel and Krauss, 1998) and has widespread impact on metabolites (Dong et al., 2023), but its impact on CHD through metabolites remains less studied. Metabolites with more than 50% missing values were excluded from the analysis. For the remaining metabolites, missing values were imputed by assigning half of the lowest observed value (Paynter et al., 2018). Metabolomic profiles were log-transformed. We further removed an outlier by inspecting a plot of the top two principal components of metabolomic profiles. Smoking status, alcohol consumption, and education level were included as covariates, with missing values imputed using multiple imputation by chained equations (van Buuren and Groothuis-Oudshoorn, 2011). After these steps, the final dataset contained 2150 samples with 366 metabolite variables. The prevalence of disease in the population was estimated as a weighted sum of disease prevalence across ethnic groups, using data from the American Heart Association’s Heart Disease and Stroke Statistics report (Tsao et al., 2022).

### 6.1. Preliminary analysis: Evidence of weak effects

In this section, we show that the data exhibited evidence of mediation signals in the paths of interest (BMI-metabolites-CHD as in Figure 3), particularly from weak mediators. Figure 3(a) presents a conceptual diagram of the hypothesized mediation structure in the WHI case-control study, where BMI influences CHD risk through high-dimensional metabolomics profiles. As a preliminary analysis, we first conducted exposure-mediator association tests to identify a subset of mediator candidates, using the significance threshold of 0.05 for p-values. Figure 3(b) displays a histogram of p-values for testing the effect of the subset of mediators on the outcome, conditional on BMI. Both the exposure–mediator and mediator–outcome tests were marginal tests and were adjusted for relevant covariates including smoking status, alcohol consumption, and education level. The red dashed line represents the estimated null density level, computed as the proportion of null mediators using Storey’s method (Storey, 2002), multiplied by the expected uniform density under the null hypothesis. P-values above this level are more likely to originate from mediators with a non-zero effect on the outcome. The prevalence of such p-values, particularly for *p >* 0.05, suggests the presence of diffuse weak mediation signals. This pattern contrasts with the expected distribution under a sparse alternative, where only a few strong mediators produce a sharp concentration of p-values near zero, while the rest follow a uniform distribution under the null.

**Fig 3.**
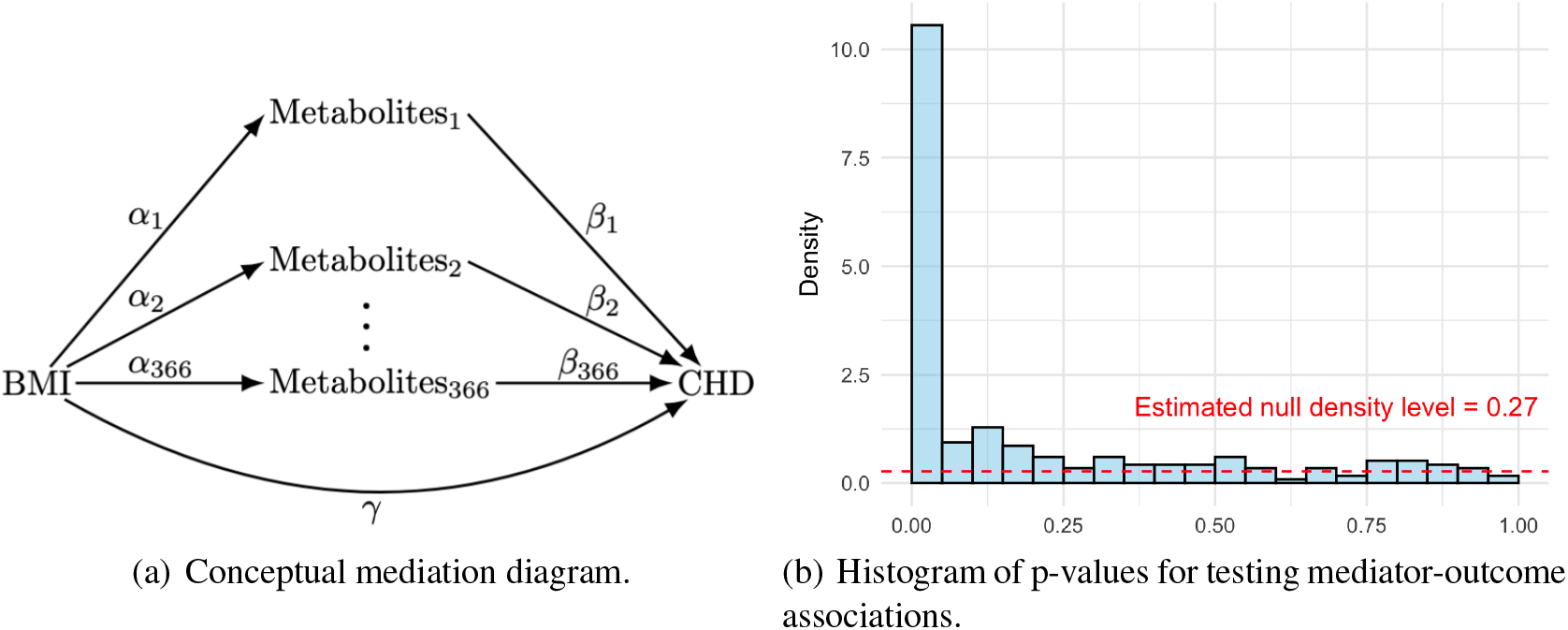
(a) Conceptual diagram illustrating the hypothesized mediation structure between BMI, metabolomics mediators, and CHD in the WHI case-control study. (b) Histogram of p-values for testing β_i_ = 0 in the outcome model, where β_i_ represents the effect of the i-th metabolite on CHD, conditional on BMI. The tests shown are restricted to mediator candidates that passed a prior test for exposure-mediator associations (i.e., with α_i_ significantly different from 0). The red dashed line represents the estimated density of null p-values using Storey’s method.

In addition, a total of 123 metabolites exhibited statistically significant associations in both marginal tests on exposure-mediator association and mediator-outcome association. However, HIMA and HDMA identified only 9 and 13 mediators, respectively, as they primarily capture mediators with strong effects. This contrast highlights the importance of capturing weaker mediation effects, which may collectively contribute to the mediation process for BMI and CHD, as well as other potential exposure-outcome relationships of interest, especially those with widespread impact on molecular phenotypes (Moore et al., 2014; Sebastiani et al., 2024).

### 6.2. Comparison and interpretation of total mediation effect estimates

Table 3 summarizes the total mediation estimates of BMI on CHD through metabolites obtained from different methods. For HIMA, HDMA, and BAMA, empirical 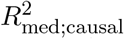 values were calculated based on the component-wise mediation estimates. The confidence intervals (CIs) for the proposed estimates were calculated using the jackknife method, where each iteration involved omitting one individual from the dataset and recomputing the estimates. The standard error of the estimator was obtained from the variability of the jackknife replicates, and 95% CIs were then calculated using a normal approximation based on the jackknife standard error. For comparison, bootstrap-based CIs were used for HIMA and HDMA, while posterior credible intervals derived from the posterior distribution were reported for BAMA. The coverage probability and computational feasibility of confidence interval was evaluated through simulation and presented in Supplementary Material Section 2.9 (Kang et al., 2025). We also evaluated the coverage probability of jackknife-based CI procedure and observed conservative yet effective nominal coverage levels under the null in simulations (Details in Supplementary Section 2.2, Kang et al. (2025)).

**Table 3.** Total mediation effect of metabolites from BMI to CHD in WHI nested case-control study. 95% confidence interval is presented in parenthesis.

| | $\hat{R}_{\text{med;causal}}^2$ | $\hat{\alpha}^\top \hat{\beta}$ | $\sum_i (\hat{\alpha}_i \hat{\beta}_i)^2$ | $\hat{Q}_{\text{med}}^2$ | $\frac{\hat{\alpha}^\top \hat{\beta}}{\hat{\alpha}^\top \hat{\beta} + \hat{\gamma}}$ |
| --- | --- | --- | --- | --- | --- |
| Proposed | 0.0076<br>(0.0033, 0.0119) | - | - | 0.8855<br>(0.5732, 1) | - |
| HIMA | 0.0033<br>(0.0019, 0.0294) | 0.09<br>(0.0376, 0.1744) | 0.0031<br>(0.0022, 0.0438) | 0.8962<br>(0.1961, 0.9989) | 0.821<br>(0.2027, 5.6155) |
| HDMA | 0.0174<br>(0.0284, 0.1506) | 0.0082<br>(-0.3889, 0.4528) | 0.0162<br>(0.0363, 0.2787) | 0.4443<br>(0.1141, 0.9985) | 0.0517<br>(-24.5906, 15.0723) |
| BAMA | 0.0002<br>(0.0001, 0.0002) | 0.0231<br>(0.0122, 0.0344) | 0.0002<br>(0.0001, 0.0002) | 0.4954<br>(0.0873, 0.9947) | 0.7361<br>(0.02538, 2.0473) |

Notably, evidence of opposing component-wise mediation effect estimates was observed across methods. HDMA method produced the smallest POC mediation effect estimate 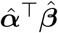. Among the 13 significant mediators identified by HDMA, 7 exhibited positive effects while 6 exhibited negative effects, leading to substantial cancellation in the total estimate. In contrast, HIMA identified 9 significant mediators, 8 of which had positive effects, resulting in a POC mediation effect estimate that was approximately ten times larger than that of HDMA, despite having a much smaller 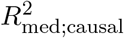 estimate. While HDMA had the largest 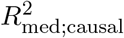 estimate, it included three potential non-mediators with p-values of *α >* 0.05, i.e., these three non-mediators may be included due to an inflated type I error in estimating *α*, which arose from not accounting for case-control selection.

In contrast, our proposed method estimated the total mediation effect of BMI on CHD through the selected set of 366 metabolites as 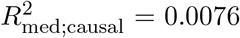, with a 95% CI of (0.0033, 0.0119), indicating that approximately 0.76% of the variance in the underlying liability for CHD was causally explained by the metabolomic mediation pathway. Although the estimated 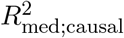 value may appear small, it reflects a statistically significant total mediation effect of a single exposure on disease liability through thousands of high-dimensional metabolomic features. In fact, small absolute mediation effects are expected herein, given the limited total effect of BMI on CHD liability estimated as 0.0086. On the other hand, the proportion of the exposure’s explained variance on the liability scale that is accounted for by metabolomic profiles, as quantified by 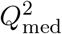, was estimated at 88.55%, with a 95% CI of (57.32%, 100%), suggesting that a substantial portion of BMI’s effect on CHD liability was mediated through these profiles.

In high-dimensional omics studies, small 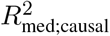 values on an absolute scale can reflect meaningful biological signal. Unlike existing approaches that focus on selecting mediators with relatively strong individual effects (e.g., HIMA and HDMA), our method aggregates contributions across weak mediators, yielding a 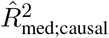 that almost doubled the value reported by HIMA/much higher than BAMA, capturing the total mediation effect that is missed by existing methods (Table 3).

## 7. Discussion

We have developed a novel variance-based total mediation effect measure for binary outcomes and high-dimensional mediators. Existing mediation measures for binary outcomes are typically mean-based and the choice of the measure often varies by disease prevalence. The key contributions of our work include (1) a new variance-based measure designed for binary outcomes, which addresses the challenge of bidirectional componentwise effects, invariant to disease prevalence, and provides a clear causal interpretation; (2) an efficient estimation procedure that accounts for ascertainment bias and effectively handles non-mediators and weak mediators. Many exposures, such as age, alcohol consumption, and BMI, have widespread effects on omics variables and may impact health outcomes through such omics mediators. The new measure and estimation method, motivated by real data, address the critical gaps in handling these broadly impactful exposures and realistic mediation patterns. To ensure theoretical validity, we demonstrate that under mild conditions, and without requiring exact mediator selection, which permits the inclusion of mediators with weak effects, our estimators converge reliably when the false discovery rate is appropriately controlled (see Section 1 of the Supplementary Material (Kang et al., 2025)). Through simulation studies, we demonstrated its robustness in estimating mediation effects under varying scenarios, and its applicability was further validated in the WHI study. The proposed method is implemented in the R package “r2MedCausal”, available on GitHub at https://github.com/zhiyu-kang/r2MedCausal.

Despite its strengths, our proposed 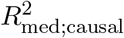 estimation procedure has some limitations. First of all, our estimation method specifically targets capturing weak but non-negligible mediation effects. It does not take into account strong effects explicitly. In other words, when only a few (≤ 5) sparse strong effects exist, existing high-dimensional mediation analysis methods with POC measures are more effective (Details in Supplementary Section 2.4, Kang et al. (2025)). Therefore, in practice, we recommend performing preliminary analysis as in Section 5.1 before using our method. Employing a multiple-step procedure may solve this issue (Li et al., 2025); however, it is not directly assessed in this work. Secondly, while PCGC was proposed to account for ascertainment bias in case-control studies, it may be less efficient than likelihood-based methods for cohort studies. We established its theoretical property under conditional parallel mediator assumption; however, we showed that our method was fairly robust to mild violation of this assumption (Supplementary Section 2.8). We recommend checking the residual distribution and consider pre-screening, increasing the number of principal components, or other batch effect correction methods if strong dependencies are observed. Thirdly, as a proportion estimator, 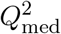 becomes unstable when the denominator gets close to zero (e.g., total effect *<* 0.003; see Supplementary Section 2.6). In practice, we suggest pursuing mediation analysis only when sufficiently large total effect is observed, following Baron and Kenny (1986). Lastly, we used jackknife for CI estimation, which we have refined its computation in our R package; however, it could still be time-consuming with large sample sizes (Supplementary Section 2.9). In terms of further research directions, we are interested in extending our causal mediation framework to multi-omics or tensor data, where mediators originate from multiple sources on different scales.

## Acknowledgments

Z. Kang and L. Chen contributed equally. C. Li and T. Yang jointly supervised the project. Correspondence: C. Li and T. Yang. T. Yang would like to thank Josey Sorenson for helping improve the interpretation of the proposed measures. The WHI program is funded by the National Heart, Lung, and Blood Institute, National Institutes of Health, U.S. Department of Health and Human Services through contracts HHSN268201600018C, HHSN268201600001C, HHSN268201600002C, HHSN268201600003C, and HHSN268201600004C. Metabolomic analyses were funded by NHLBI contract HHSN268201300008C. This manuscript was not prepared in collaboration with investigators of the WHI, has not been reviewed and/or approved by WHI, and does not necessarily reflect the opinions of the WHI investigators or the NHLBI. The WHI dataset used for the analyses described in this manuscript was obtained from dbGaP at https://www.ncbi.nlm.nih.gov/gap through accession number phs001334. The authors acknowledge the Minnesota Supercomputing Institute (MSI) at the University of Minnesota for providing resources that contributed to the research results reported in this paper. The authors also acknowledge the use of generative AI tools to assist in refining sentence structure, improving clarity, and ensuring conciseness in the manuscript. The authors would like to thank the referee, the Associate Editor and the Editor for the constructive feedback which greatly improved this work.

## Significance Statement

Modern biomedical studies often measure thousands of molecular features to understand how exposures influence disease risk. However, existing measures of total mediation are not well suited for high-dimensional settings, especially when many small effects act in opposite directions. We propose a new causally interpretable measure that summarizes the overall mediation effect for binary disease outcomes, together with a fast estimation procedure that applicable to case–control sampling and large numbers of mediators. An application to a metabolomics study demonstrates that the proposed method can detect mediation signals missed by existing approaches. This work provides a practical tool for studying mediation effects in high-dimensional omics data.

## Supplementary Material

### Technical proofs

Contains all formal proofs of the main theoretical results and supporting lemmas, together with the full list of technical conditions used throughout the manuscript.

### Additional simulation results

Presents extra numerical experiments evaluating estimator performance under a variety of settings.

### PCGC algorithm

Gives a step-by-step description of the PCGC regression procedure when covariates are present.

### Illustration of cross-fitting procedure

Provides a schematic diagram of the two-fold sample-splitting and cross-fitting steps.

### r2MedCausal

Provides the R package “r2MedCausal” implementing the proposed method described in this manuscript. The package is also publicly available at https://github.com/zhiyu-kang/r2MedCausal.

