## Supplementary material for "Estimating total mediation effects of high-dimensional omics mediators in case-control studies": Supp files

### SUPPLEMENT TO "ESTIMATION OF TOTAL MEDIATION EFFECT FOR A BINARY TRAIT IN A CASE-CONTROL STUDY FOR HIGH-DIMENSIONAL OMICS MEDIATORS"

BY ZHIYU KANG[1], LI CHEN[2], PENG WEI[3], ZHICHAO XU[3], CHUNLIN LI[4,b], AND TIANZHONG YANG[1,a]

[1]*Division of Biostatistics and Health Data Science, University of Minnesota, Twin Cities,* [a]**

[2]*School of Statistics, University of Minnesota, Twin Cities*

[3]*Department of Biostatistics, University of Texas MD Anderson Cancer Center*

[4]*Department of Statistics, Iowa State University,* [b]**

#### 1. Technical proofs.

In this section, we prove the theoretical result in the main text.

##### 1.1. *Technical conditions.*

ASSUMPTION 1 (No Unmeasured Confounding). *The following assumptions are required for valid causal mediation analysis:*
***A1*** *No unmeasured confounding of the exposure-mediator effect.*
***A2*** *No unmeasured confounding of the mediator-outcome effect.*
***A3*** *No unmeasured confounding of the exposure-outcome effect.*
***A4*** *No mediator-outcome confounder that is itself affected by the exposure.*

ASSUMPTION 2 (Conditional Parallel Mediator Assumption). *There exist random variables* $\boldsymbol{U}$ *that have a global influence on* $\boldsymbol{M}$*, where* $M_k \perp\!\!\!\perp M_j | X, \boldsymbol{U}, \boldsymbol{C}$ *for any* $k, j \in \{1, \ldots, p\}$.

In the following discussion, we note that our proposed procedure is still valid and robust even if Assumption 2 is violated.

**The proposed measure is causally meaningful when dependencies among mediators are present.** Suppose we have the following underlying true model:

$$\boldsymbol{M} = \mathbf{A}^\top \boldsymbol{M} + \boldsymbol{\alpha}' X + \boldsymbol{\Psi}' \boldsymbol{C} + \boldsymbol{\xi}',$$
$$l = \gamma X + \boldsymbol{\beta}^\top \boldsymbol{M} + \boldsymbol{\theta}^\top \boldsymbol{C} + \epsilon,$$
$$Y = \mathbf{1}(l > t),$$

where $\mathbf{A}$ encodes a DAG with $\mathbf{A}_{k,j} \neq 0$ if and only if there exists an edge from $\boldsymbol{M}_k$ to $\boldsymbol{M}_j$. Define

$$\alpha_j = \sum_{k \in \mathrm{AN}(j)} \mathbf{B}_{k,j} \alpha'_k + \alpha'_j, \quad \boldsymbol{\xi}_j = \sum_{k \in \mathrm{AN}(j)} \mathbf{B}_{k,j} \boldsymbol{\xi}'_k + \boldsymbol{\xi}'_j, \text{ and } \boldsymbol{\Psi}_{j,l} = \sum_{k \in \mathrm{AN}(j)} \mathbf{B}_{k,j} \boldsymbol{\Psi}'_{k,l} + \boldsymbol{\Psi}'_{j,l},$$

where $\mathbf{B} = \mathbf{A} + \cdots + \mathbf{A}^{p+1}$. Then the underlying true model can be transformed into

$$\boldsymbol{M} = \boldsymbol{\alpha} X + \boldsymbol{\Psi} \boldsymbol{C} + \boldsymbol{\xi},$$
$$l = \gamma X + \boldsymbol{\beta}^\top \boldsymbol{M} + \boldsymbol{\theta}^\top \boldsymbol{C} + \epsilon,$$
$$Y = \mathbf{1}(l > t),$$

*Zhiyu Kang and Li Chen contributed equally to this work. Chunlin Li and Tianzhong Yang jointly supervised the project. Corresponding authors: Chunlin Li and Tianzhong Yang.

which corresponds to the structural equation model we use, except that $\boldsymbol{\xi}$ may exhibit cross-component correlation. If we assume that $\mathbb{P}(\alpha_k \neq 0) = \pi_\alpha$, $\mathbb{E}[\alpha_k] = 0$ and $\mathrm{var}[\alpha_k \mid \alpha_k \neq 0] = \sigma_\alpha^2$, then for the proposed measure, it still holds that

$$R^2_{\text{med;causal}} = \sigma_\alpha^2 \sigma_{11}^2 \mathrm{var}(X)/\mathrm{var}_{\boldsymbol{\alpha},\boldsymbol{\beta},l}(l),$$

and

$$Q^2_{\text{med}} = \frac{\sigma_\alpha^2 \sigma_{11}^2}{\sigma_\alpha^2 \sigma_{11}^2 + \gamma^2}.$$

**The proposed estimation procedure is robust against sparse correlations.** In Section 4.1, we need to estimate $\mathrm{var}[\boldsymbol{\alpha}]$, and in Section 4.3, we need to apply PCGC regression with $\hat{\boldsymbol{\xi}}_{\hat{\mathcal{T}}}$ as the standardized design matrix to estimate $\hat{\sigma}_{11}^2$. Accordingly, provided that these two estimation steps retain the required convergence properties, which is plausible when mediator causal dependencies are relatively sparse and of moderate strength, our procedure should remain robust even when $\boldsymbol{\xi}$ exhibits cross-component correlation. Please refer to the additional simulation study (Supplementary Section 2.8) for a numerical evaluation of the robustness of the proposed method under increasing levels of residual correlation among mediators.

ASSUMPTION 3. *Let $\widetilde{\mathbf{X}}_i = (I_p \otimes X_i, I_p \otimes \boldsymbol{C}_i^\top)$; there exists a positive constant $R$ such that*

$$0 < \frac{1}{R} < \lambda_{\min}\Big(\frac{1}{n}\sum_{i=1}^n \widetilde{\mathbf{X}}_i^\top \widetilde{\mathbf{X}}_i\Big) \leq \lambda_{\max}\Big(\frac{1}{n}\sum_{i=1}^n \widetilde{\mathbf{X}}_i^\top \widetilde{\mathbf{X}}_i\Big) < R < \infty,$$

*and*

$$0 < \frac{1}{R} < \pi_\alpha \sigma_\alpha^2 < R < \infty,$$

*and for all $i$,*

$$0 < \frac{1}{R} < w_i < R < \infty.$$

ASSUMPTION 4. *We have $\mathbf{E}[\xi_{ij}^4] < \infty$ and $\sup_{jk} \mathbf{E}[(\xi_{ij}\xi_{ik})^4] < \infty$ for $i = 1, \ldots, n$ and $j, k = 1, \ldots, p$.*

Assumptions 3 and 4 are standard conditions in case-control studies, see Conditions (C2),(C3) and (C9) in Sofer et al. (2017) for an example.

ASSUMPTION 5. *There exists a constant $c > 0$, such that $\mathbb{E}[\mathrm{var}(X \mid \boldsymbol{M})] > c$.*

ASSUMPTION 6. *Let $\boldsymbol{c}_i = (X_i, \boldsymbol{C}_i, \boldsymbol{W}_i)^\top$ and $\widehat{\boldsymbol{c}}_i = (X_i, \boldsymbol{C}_i, \widehat{\boldsymbol{W}}_i)^\top$. Assume that there exists a constant $C > 0$, such that $\max(\|\boldsymbol{c}_i\|_2, \|\widehat{\boldsymbol{c}}_i\|_2) < C$ and $\frac{1}{n}\sum_{i=1}^n \|\widehat{\boldsymbol{c}}_i - \boldsymbol{c}_i\|_2 = o_p(1)$.*

Assumption 5 ensures that $\gamma$ is identifiable and Assumption 6 is very mild in the factor model literature, with Theorem 4 in Fan, Liao and Mincheva (2013) as an example.

Following the derivation of heritability estimation with known fixed effects in Golan, Lander and Rosset (2014), we have

$$\mathbb{E}\left[Z_{ij} \mid \mathcal{S} = 1; G_{ij}, t_i, t_j\right] = \frac{A'(0; t_i, t_j)}{B(0; t_i, t_j)} G_{ij} + b_{ij}(G_{ij}; t_i, t_j) G_{ij}^2,$$

where

$$A\left(G_{ij};t_i,t_j\right)=\frac{\left(1-P_i\right)\left(1-P_j\right)}{\sqrt{P_i\left(1-P_i\right)}\sqrt{P_j\left(1-P_j\right)}}\mathbb{P}\left(y_i=y_j=1;G_{ij},t_i,t_j\right)+$$
$$\frac{K(1-P)}{P(1-K)}\frac{-P_i\left(1-P_j\right)}{\sqrt{P_i\left(1-P_i\right)}\sqrt{P_j\left(1-P_j\right)}}\mathbb{P}\left(y_i=0,y_j=1;G_{ij},t_i,t_j\right)+$$
$$\frac{K(1-P)}{P(1-K)}\frac{-P_j\left(1-P_i\right)}{\sqrt{P_i\left(1-P_i\right)}\sqrt{P_j\left(1-P_j\right)}}\mathbb{P}\left(y_i=1,y_j=0;G_{ij},t_i,t_j\right)+$$
$$\left(\frac{K(1-P)}{P(1-K)}\right)^2\frac{P_iP_j}{\sqrt{P_i\left(1-P_i\right)}\sqrt{P_j\left(1-P_j\right)}}\mathbb{P}\left(y_i=0,y_j=0;G_{ij},t_i,t_j\right),$$

and

$$B\left(\rho;t_i,t_j\right)=\mathbb{P}(\mathcal{S}=1;\rho,t_i,t_j).$$

Denote $\Delta_{ij}=\frac{A'(0;t_i,t_j)}{B(0;t_i,t_j)\sigma_{11}^2}=\frac{\varphi(t_i)\varphi(t_j)\left[1-(P_i+P_j)\left(\frac{P-K}{P(1-K)}\right)+P_iP_j\left(\frac{P-K}{P(1-K)}\right)^2\right]}{\sqrt{P_i(1-P_i)}\sqrt{P_j(1-P_j)}\left(K_i+(1-K_i)\frac{K(1-P)}{P(1-K)}\right)\left(K_j+(1-K_j)\frac{K(1-P)}{P(1-K)}\right)}$.

ASSUMPTION 7. *Let* $e_{ij}=Z_{ij}-\mathbb{E}\left[Z_{ij}\mid\mathcal{S}=1;G_{ij},t_i,t_j\right]$, *we assume that*

$$\frac{\sum_{i<j}\Delta_{ij}b_{ij}(G_{ij})G_{ij}^3}{\sum_{i<j}\Delta_{ij}^2G_{ij}^2}\to 0,$$

*and*

$$\frac{\sum_{i<j}\Delta_{ij}G_{ij}e_{ij}}{\sum_{i<j}\Delta_{ij}^2G_{ij}^2}\to 0,$$

*as* $n\to\infty$.

The consistency of the PCGC estimator without fixed effects when $\frac{p}{n}$ converges to a positive constant was established in Bonnet (2018). Under their assumptions, the first part of Assumption 7 is implied by their Lemma 1 and Lemma 2, and the second part of Assumption 7 is implied by their Lemma 4.

ASSUMPTION 8. *We have that*

$$\max\Big(\sum_{i<j}|\Delta_{ij}G_{ij}-\widehat{\Delta}_{ij}\widehat{G}_{ij}|,|\sum_{i<j}(\widehat{\Delta}_{ij}^2\widehat{G}_{ij}^2-\Delta_{ij}^2G_{ij}^2)|\Big)=o_p(n(n-1)).$$

ASSUMPTION 9. *There exists a positive constant* $0<r<0.5$ *such that*

$$0<r<P<1-r<1,0<r<K<1-r<1,$$

*and for all* $i$,

$$0<r<P_i<1-r<1,0<r<K_i<1-r<1.$$

For $\sigma>0$, let

$$L_n(\sigma)=\frac{1}{n(n-1)}\sum_{i<j}\left(Z_{ij}-\sigma^2\widehat{\Delta}_{ij}\widehat{G}_{ij}\right)^2,$$

and

$$G_n(\sigma) = \frac{1}{n(n-1)} \sum_{i<j} \left( \widetilde{Z}_{ij} - \sigma^2 \Delta_{ij} G_{ij} \right)^2,$$

where $\widetilde{Z}_{ij} = Z_{ij} - \frac{\sum_{i<j} \Delta_{ij} b_{ij}(G_{ij}) G_{ij}^3}{\sum_{i<j} \Delta_{ij}^2 G_{ij}^2} \Delta_{ij} G_{ij}$.

ASSUMPTION 10. *There exists a function $G(\sigma)$ possessing a unique minimizer such that $\forall \sigma > 0$, $G_n(\sigma) \to G(\sigma)$ in probability as $n \to \infty$.*

Assumption 10 is an assumption of convenience, and relaxation could be employed by studying sub-sequences of $G_n(\cdot)$, see Theorem 10.8 and 10.9 in Rockafellar (1970). A similar condition was adopted in Condition (C1) of Stefanski and Carroll (1985) to prove the consistency of the estimator in logistic regression with measurement error.

1.2. *Proof of Theorem 1.* This is a direct result of Lemma 1.1 (see below) and the Bolzano-Weierstrass theorem.

1.3. *Technical lemmas.* Given the confusion matrix, the variance to be estimated by the PCGC model is the variance of the random effect:

$$\text{TP1} \times \frac{\sigma_{11}^2}{p\pi_{11}} + \text{FP1} \times \frac{\sigma_{01}^2}{p\pi_{01}},$$

where TP1 and FP1 represent the number of $\boldsymbol{M}_{\mathcal{T}}$ and $\boldsymbol{M}_{\mathcal{I}_1}$ that are not filtered out, respectively. Furthermore, we let FN1 and TN1 represent the numbers of $\boldsymbol{M}_{\mathcal{T}}$ and $\boldsymbol{M}_{\mathcal{I}_1}$ that are filtered out, respectively.

ASSUMPTION 11. *There exist constants $c_{\text{TP}}, c_{\text{TN}} > 0$ and $c_{\text{FP}}, c_{\text{FN}} \geq 0$ such that $c_{\text{TP}} + c_{\text{FP}} + c_{\text{TN}} + c_{\text{FN}} = 1$ and*

$$\lim_{p\to\infty, n\to\infty} \text{TP} : \text{FP} : \text{TN} : \text{FN} = c_{\text{TP}} : c_{\text{FP}} : c_{\text{TN}} : c_{\text{FN}}.$$

LEMMA 1.1. *Under Assumptions in Lemma 1.2, Lemma 1.3, and Lemma 1.4, for any $\epsilon > 0$, there exists $\delta > 0$, such that if $\frac{c_{\text{TP}}}{c_{\text{TP}} + c_{\text{FP}}} > 1 - \delta$, we have*

$$\lim_{p\to\infty, n\to\infty} \mathbb{P}(|\widehat{R}^2_{\text{med;causal}} - R^2_{\text{med;causal}}| < \epsilon) = 1,$$

*and*

$$\lim_{p\to\infty, n\to\infty} \mathbb{P}(|\widehat{Q}^2_{\text{med}} - Q^2_{\text{med}}| < \epsilon) = 1.$$

PROOF. By the law of large number, under Assumption 11, we have

$$\lim_{p\to\infty, n\to\infty} \frac{\text{TP1}}{\text{TP1} + \text{FN1}} = \frac{c_{\text{TP}}}{c_{\text{TP}} + c_{\text{FN}}}, \qquad \lim_{p\to\infty, n\to\infty} \frac{\text{FP1}}{\text{FP1} + \text{TN1}} = \frac{c_{\text{FP}}}{c_{\text{FP}} + c_{\text{TN}}}.$$

Regarding the estimate of $\sigma_\alpha^2$, we have $\text{var}(\alpha) = \pi_\alpha \sigma_\alpha^2$, where $\pi_\alpha = c_{\text{TP}} + c_{\text{FN}}$. Given that $\hat{\pi}_\alpha = c_{\text{TP}} + c_{\text{FP}}$ we can have the estimate

$$\hat{\sigma}_\alpha^2 = \frac{\widehat{\text{var}(\alpha)}}{\hat{\pi}_\alpha} = \frac{\widehat{\text{var}(\alpha)}}{c_{\text{TP}} + c_{\text{FP}}} = \frac{\widehat{\text{var}(\alpha)}}{c_{\text{TP}} + c_{\text{FN}}} \frac{c_{\text{TP}} + c_{\text{FN}}}{c_{\text{TP}} + c_{\text{FP}}} = \sigma_\alpha^2 \frac{\widehat{\text{var}(\alpha)}}{\text{var}(\alpha)} \frac{c_{\text{TP}} + c_{\text{FN}}}{c_{\text{TP}} + c_{\text{FP}}}.$$

By Lemma 1.2 and Lemma 1.4, we have:

$$\hat{\sigma}_\alpha^2 \times \hat{\sigma}_{11}^2 \to \sigma_\alpha^2 \frac{c_{\mathrm{TP}} + c_{\mathrm{FN}}}{c_{\mathrm{TP}} + c_{\mathrm{FP}}} \times \left( \frac{c_{\mathrm{TP}}}{c_{\mathrm{TP}} + c_{\mathrm{FN}}} \sigma_{11}^2 + \frac{c_{\mathrm{FP}}}{c_{\mathrm{FP}} + c_{\mathrm{TN}}} \sigma_{01}^2 \right)$$
$$= \sigma_\alpha^2 \times \left( \frac{c_{\mathrm{TP}}}{c_{\mathrm{TP}} + c_{\mathrm{FP}}} \sigma_{11}^2 + \frac{c_{\mathrm{FP}}}{c_{\mathrm{TP}} + c_{\mathrm{FP}}} \frac{\pi_\alpha}{1-\pi_\alpha} \sigma_{01}^2 \right)$$

as $p, n \to \infty$. The above arguments along with Assumption 3 and Lemma 1.3 complete the proof. Specifically, $\delta$ can be chosen to be

$$\min\left(\frac{\epsilon\,\mathrm{var}(l)}{2\sigma_x^2\sigma_\alpha^2|\sigma_{01}^2\frac{\pi_\alpha}{1-\pi_\alpha} - \sigma_{11}^2|}, \frac{\epsilon(\sigma_\alpha^2\sigma_{11}^2+\gamma^2)^2}{4|\sigma_{01}^2\frac{\pi_\alpha}{1-\pi_\alpha} - \sigma_{11}^2|(2\gamma^2 + \epsilon(\sigma_\alpha^2\sigma_{11}^2+\gamma^2))}, 1\right),$$

if $\sigma_{11}^2 \neq \sigma_{01}^2 \times \frac{\pi_\alpha}{1-\pi_\alpha}$ or $\delta = 1$ if $\sigma_{11}^2 = \sigma_{01}^2 \times \frac{\pi_\alpha}{1-\pi_\alpha}$. □

LEMMA 1.2. *Under Assumption 1, 2, 3 and 4, we have*

$$|\widehat{\mathrm{var}(\alpha)} - \mathrm{var}(\alpha)| = O_p\left(\frac{1}{\sqrt{p}} + \frac{1}{\sqrt{n}}\right).$$

PROOF. Let $\widetilde{\alpha} = (\mathrm{Vec}(\alpha)^\top, \mathrm{Vec}(\Psi^\top))^\top$ and $D(\boldsymbol{u}) = Q(\widetilde{\alpha}^* + \boldsymbol{u}) - Q(\widetilde{\alpha}^*) = J_1(\boldsymbol{u}) + J_2(\boldsymbol{u})$, where $Q$ is the quadratic loss and

$$J_1(\boldsymbol{u}) = -2\left(\sum_{i=1}^n w_i \boldsymbol{\xi}_i^\top \widetilde{\mathbf{X}}_i\right)\boldsymbol{u},$$
$$J_2(\boldsymbol{u}) = \boldsymbol{u}^T\left(\sum_{i=1}^n w_i \widetilde{\mathbf{X}}_i^\top \widetilde{\mathbf{X}}_i\right)\boldsymbol{u}.$$

Let $\widetilde{\mathbf{W}} = \mathrm{diag}(w_1, \ldots, w_n) \otimes I_p$, $\widetilde{\boldsymbol{\xi}} = (\boldsymbol{\xi}_1^\top, \ldots, \boldsymbol{\xi}_n^\top)^\top$ and $\widetilde{\mathbf{X}} = (\widetilde{\mathbf{X}}_1^\top, \ldots, \widetilde{\mathbf{X}}_n^\top)^\top$, where $\mathrm{diag}(w_1, \ldots, w_n)$ is the $n \times n$ diagonal matrix with the $(w_1, \ldots, w_n)$ on its main diagonal and $\otimes$ denotes the Kronecker product.

By Lemma 1.5, we have

$$|J_1(\boldsymbol{u})| = 2\left\langle \widetilde{\mathbf{X}}^\top \widetilde{\mathbf{W}} \widetilde{\boldsymbol{\xi}}, \boldsymbol{u}\right\rangle \le 2\|\widetilde{\mathbf{X}}^\top \widetilde{\mathbf{W}} \widetilde{\boldsymbol{\xi}}\|\|\boldsymbol{u}\| = 2O_p(\sqrt{np})\|\boldsymbol{u}\|.$$

By Assumption 3, we have

$$J_2(\boldsymbol{u}) \ge n\|\boldsymbol{u}\|^2 R^{-2}.$$

Then for any $0 < c < 1$, we can choose sufficiently large $M$ such that

$$\mathbb{P}\left(\inf_{\|\boldsymbol{u}\|=1} Q(\widetilde{\alpha}^* + Mp/\sqrt{n}\boldsymbol{u}) - Q(\widetilde{\alpha}^*) > 0\right) \ge 1 - c$$

for $n$ sufficiently large. Thus $\|\widehat{\widetilde{\alpha}^*} - \widetilde{\alpha}^*\| = O_p(\frac{p}{\sqrt{n}})$. Then we have

$$\left|\frac{1}{p}\|\widehat{\boldsymbol{\alpha}}\|^2 - \frac{1}{p}\|\boldsymbol{\alpha}^*\|^2\right| = O_p\left(\frac{1}{\sqrt{n}}\right).$$

By the fact that $|\,\mathrm{var}(\alpha) - \frac{1}{p}\|\boldsymbol{\alpha}^*\|^2| = O_p(\frac{1}{\sqrt{p}})$, we have

$$|\widehat{\mathrm{var}(\alpha)} - \mathrm{var}(\alpha)| = O_p\left(\frac{1}{\sqrt{p}} + \frac{1}{\sqrt{n}}\right).$$

□

LEMMA 1.3. *Under Assumptions 1, 2, 3, 5 and 6, we have* $\|\widehat{\boldsymbol{\nu}} - \boldsymbol{\nu}_0\|_2 = o_p(1)$, *where* $\boldsymbol{\nu}$ *corresponds to* $(X_i, \boldsymbol{C}_i, \boldsymbol{W}_i)$.

PROOF. Let

$$L_n(\boldsymbol{\nu}) = \frac{1}{n}\sum_1^n w_i \left\{ y_i \log \Phi\left(\widehat{\boldsymbol{c}}_i^\top \boldsymbol{\nu}\right) + (1-y_i)\log\Phi\left(-\widehat{\boldsymbol{c}}_i^\top \boldsymbol{\nu}\right)\right\}$$

and $\widehat{\boldsymbol{\nu}} = \arg\max_{\boldsymbol{\nu}} L_n(\boldsymbol{\nu})$. Let

$$G_n(\boldsymbol{\nu}) = \frac{1}{n}\sum_1^n w_i \left\{ \Phi\left(\boldsymbol{c}_i^\top \boldsymbol{\nu}_0\right)\log\Phi\left(\boldsymbol{c}_i^\top \boldsymbol{\nu}\right) + \Phi\left(-\boldsymbol{c}_i^\top \boldsymbol{\nu}_0\right)\log\Phi\left(-\boldsymbol{c}_i^\top \boldsymbol{\nu}\right)\right\}.$$

Then

$$L_n(\boldsymbol{\nu}) - G_n(\boldsymbol{\nu}) = R_{n,1}(\boldsymbol{\nu}) + R_{n,2}(\boldsymbol{\nu}),$$

where

$$R_{n,1}(\boldsymbol{\nu}) = \frac{1}{n}\sum_1^n w_i \left\{ \left(y_i - \Phi\left(\boldsymbol{c}_i^\top \boldsymbol{\nu}_0\right)\right)\log\Phi\left(\widehat{\boldsymbol{c}}_i^\top \boldsymbol{\nu}\right) + \left(1 - y_i - \Phi\left(-\boldsymbol{c}_i^\top \boldsymbol{\nu}_0\right)\right)\log\Phi\left(-\widehat{\boldsymbol{c}}_i^\top \boldsymbol{\nu}\right)\right\},$$

and

$$\begin{aligned} R_{n,2}(\boldsymbol{\nu}) =& \frac{1}{n}\sum_1^n w_i \{\Phi\left(\boldsymbol{c}_i^\top \boldsymbol{\nu}_0\right)\left(\log\Phi\left(\boldsymbol{c}_i^\top \boldsymbol{\nu}\right) - \log\Phi\left(\widehat{\boldsymbol{c}}_i^\top \boldsymbol{\nu}\right)\right) \\ &+ \Phi\left(-\boldsymbol{c}_i^\top \boldsymbol{\nu}_0\right)\left(\log\Phi\left(-\boldsymbol{c}_i^\top \boldsymbol{\nu}\right) - \log\Phi\left(-\widehat{\boldsymbol{c}}_i^\top \boldsymbol{\nu}\right)\right)\}. \end{aligned}$$

For any fixed $\boldsymbol{\nu}$, under Assumption 3 and 6, $R_{n,1}(\boldsymbol{\nu})$ has mean zero and asymptotically negligible variance, and

$$|R_{n,2}(\boldsymbol{\nu})| \le 2R\frac{1}{n}\sum_{i=1}^n \frac{\|\widehat{\boldsymbol{c}}_i - \boldsymbol{c}_i\|_2 \|\boldsymbol{\nu}\|_2}{\sqrt{2\pi}\Phi(-C\|\boldsymbol{\nu}\|_2)} = o_p(1).$$

Note $G_n(\boldsymbol{\nu})$ converges in probability to $\mathbb{E}[G_n(\boldsymbol{\nu})]$ for any fixed $\boldsymbol{\nu}$ and $\mathbb{E}[G_n(\boldsymbol{\nu})]$ has a unique maximum at $\boldsymbol{\nu}_0$ under Assumption 5. We have that $L_n(\boldsymbol{\nu})$ converges in probability to $\mathbb{E}[G_n(\boldsymbol{\nu})]$ for any fixed $\boldsymbol{\nu}$. This completes the proof by Theorem 2.2 in Andersen and Gill (1982). □

We define $\sigma^2_{11(\text{adj})} = \frac{c_{\text{TP}}}{c_{\text{TP}}+c_{\text{FN}}}\sigma^2_{11} + \frac{c_{\text{FP}}}{c_{\text{FP}}+c_{\text{TN}}}\sigma^2_{01}$ and introduce the following lemma.

LEMMA 1.4. *Under Assumption 1, 2, 7, 8, 9, 10 and 11*

$$|\hat\sigma^2_{11} - \sigma^2_{11(\text{adj})}| = o_p(1).$$

PROOF. Note that

$$\begin{aligned} L_n(\sigma) - G_n(\sigma) =& \frac{2\sigma^2}{n(n-1)}\sum_{i<j}(\Delta_{ij}G_{ij} - \widehat{\Delta}_{ij}\widehat{G}_{ij})Z_{ij} \\ &+ \frac{\sigma^4}{n(n-1)}\sum_{i<j}(\widehat{\Delta}^2_{ij}\widehat{G}^2_{ij} - \Delta^2_{ij}G^2_{ij}) \end{aligned}$$

$$- \frac{2\sigma^2}{n(n-1)} \frac{\sum_{i<j} \Delta_{ij} b_{ij}(G_{ij}) G_{ij}^3}{\sum_{i<j} \Delta_{ij}^2 G_{ij}^2} \sum_{i<j} \Delta_{ij}^2 G_{ij}^2$$
$$+ \frac{1}{n(n-1)} \frac{\sum_{i<j} \Delta_{ij} b_{ij}(G_{ij}) G_{ij}^3}{\sum_{i<j} \Delta_{ij}^2 G_{ij}^2} \sum_{i<j} \left( 2Z_{ij} - \frac{\sum_{i<j} \Delta_{ij} b_{ij}(G_{ij}) G_{ij}^3}{\sum_{i<j} \Delta_{ij}^2 G_{ij}^2} \Delta_{ij} G_{ij} \right) G_{ij}.$$

By Assumption 9, there exists a constant $M_1 > 0$, such that $|Z_{ij}| < M_1$. Then for any fixed $\sigma > 0$, we have

$$\left| \frac{2\sigma^2}{n(n-1)} \sum_{i<j} (\Delta_{ij} G_{ij} - \widehat{\Delta}_{ij} \widehat{G}_{ij}) Z_{ij} \right| \le \frac{2M_1\sigma^2}{n(n-1)} \sum_{i<j} \left| \Delta_{ij} G_{ij} - \widehat{\Delta}_{ij} \widehat{G}_{ij} \right| = o_p(1),$$

and

$$\left| \frac{\sigma^4}{n(n-1)} \sum_{i<j} (\widehat{\Delta}_{ij}^2 \widehat{G}_{ij}^2 - \Delta_{ij}^2 G_{ij}^2) \right| = o_p(1)$$

by Assumption 8. There exists a constant $M_2 > 0$, such that $|\Delta_{ij}| < M_2$ by Assumption 9 and

$$\left| 2Z_{ij} - \frac{\sum_{i<j} \Delta_{ij} b_{ij}(G_{ij}) G_{ij}^3}{\sum_{i<j} \Delta_{ij}^2 G_{ij}^2} \Delta_{ij} G_{ij} \right| = O_p(1)$$

by Assumption 7. Then for any fixed $\sigma > 0$, we have that

$$\left| \frac{2\sigma^2}{n(n-1)} \frac{\sum_{i<j} \Delta_{ij} b_{ij}(G_{ij}) G_{ij}^3}{\sum_{i<j} \Delta_{ij}^2 G_{ij}^2} \sum_{i<j} \Delta_{ij}^2 G_{ij}^2 \right| \le 2M_2^2\sigma^2 \left| \frac{\sum_{i<j} \Delta_{ij} b_{ij}(G_{ij}) G_{ij}^3}{\sum_{i<j} \Delta_{ij}^2 G_{ij}^2} \right| = o_p(1),$$

and

$$\left| \frac{1}{n(n-1)} \frac{\sum_{i<j} \Delta_{ij} b_{ij}(G_{ij}) G_{ij}^3}{\sum_{i<j} \Delta_{ij}^2 G_{ij}^2} \sum_{i<j} \left( 2Z_{ij} - \frac{\sum_{i<j} \Delta_{ij} b_{ij}(G_{ij}) G_{ij}^3}{\sum_{i<j} \Delta_{ij}^2 G_{ij}^2} \Delta_{ij} G_{ij} \right) G_{ij} \right| = o_p(1).$$

Thus we have $|L_n(\sigma) - G_n(\sigma)| = o_p(1)$ for any fixed $\sigma > 0$.

Denote $\widehat{\sigma}_{L_n} = \arg\max_{\sigma>0} L_n(\sigma)$ and $\widehat{\sigma}_{G_n} = \arg\max_{\sigma>0} G_n(\sigma)$. By Assumption 10, both $L_n(\sigma)$ and $G_n(\sigma)$ converge to $G(\sigma)$ in probability for any $\sigma > 0$. Denote the unique minimizer as $\sigma_*$, by Theorem 2.2 in Andersen and Gill (1982), we have $|\widehat{\sigma}_{L_n}^2 - \sigma_*^2| = o_p(1)$ and $|\widehat{\sigma}_{G_n}^2 - \sigma_*^2| = o_p(1)$. Thus it remains to compute $\lim_{n\to\infty} \widehat{\sigma}_{G_n}^2$.

$$\begin{aligned}
\widehat{\sigma}_{G_n}^2 =& \frac{\sum_{i<j} \widetilde{Z}_{ij} \Delta_{ij} G_{ij}}{\sum_{i<j} \Delta_{ij}^2 G_{ij}^2} = \frac{\sum_{i<j} Z_{ij} \Delta_{ij} G_{ij}}{\sum_{i<j} \Delta_{ij}^2 G_{ij}^2} - \frac{\sum_{i<j} \Delta_{ij} b_{ij}(G_{ij}) G_{ij}^3}{\sum_{i<j} \Delta_{ij}^2 G_{ij}^2} \\
=& \frac{\sum_{i<j} \mathbb{E}\left[Z_{ij} \mid \mathcal{S}=1; G_{ij}, t_i, t_j\right] \Delta_{ij} G_{ij}}{\sum_{i<j} \Delta_{ij}^2 G_{ij}^2} + \frac{\sum_{i<j} e_{ij} \Delta_{ij} G_{ij}}{\sum_{i<j} \Delta_{ij}^2 G_{ij}^2} - \frac{\sum_{i<j} \Delta_{ij} b_{ij}(G_{ij}) G_{ij}^3}{\sum_{i<j} \Delta_{ij}^2 G_{ij}^2} \\
=& \sigma_{11(\text{adj})}^2 + \frac{\sum_{i<j} e_{ij} \Delta_{ij} G_{ij}}{\sum_{i<j} \Delta_{ij}^2 G_{ij}^2} \to \sigma_{11(\text{adj})}^2.
\end{aligned}$$

This completes the proof. □

LEMMA 1.5. *Let Assumptions 1, 2, 3 and 4 hold. Then*

$$\|\frac{1}{n} \widetilde{\mathbf{X}}^\top \widetilde{\mathbf{W}} \widetilde{\boldsymbol{\xi}}\| = O_p(\frac{p}{\sqrt{n}}).$$

PROOF. Note that

$$\|\frac{1}{n}\widetilde{\mathbf{X}}^\top\widetilde{\mathbf{W}}\widetilde{\boldsymbol{\xi}}\| \leq \left((q+1)p \sup_{j=1,\ldots,(q+1)p}\left(\frac{1}{n}\widetilde{\mathbf{X}}_{\cdot,j}^\top\widetilde{\mathbf{W}}\widetilde{\boldsymbol{\xi}}\right)^2\right)^{1/2}$$

$$\leq \sqrt{(q+1)p}\left(\sup_{j=1,\ldots,(q+1)p}\|\frac{1}{n}\widetilde{\mathbf{W}}\widetilde{\mathbf{X}}_{\cdot,j}\|^2 \sup_{\|\boldsymbol{u}\|=1}|\boldsymbol{u}^\top\widetilde{\boldsymbol{\xi}}|^2\right)^{1/2}.$$

For any $j = 1, \ldots, (q+1)p$, by Assumption 3, we have

$$\|\frac{1}{n}\widetilde{\mathbf{W}}\widetilde{\mathbf{X}}_{\cdot,j}\|^2 \leq \frac{1}{n}R^3.$$

This leads to

$$\|\frac{1}{n}\widetilde{\mathbf{X}}^\top\widetilde{\mathbf{W}}\widetilde{\boldsymbol{\xi}}\| \leq \sqrt{\frac{(q+1)p}{n}}R^{3/2}(\sup_{\|\boldsymbol{u}\|=1}|\boldsymbol{u}^\top\widetilde{\boldsymbol{\xi}}|^2)^{1/2}.$$

For sequences $l_n$ such that $l_n/\frac{(q+1)p}{\sqrt{n}} \to \infty$, there exists $K > 0$ such that

$$\mathbb{P}(\|\frac{1}{n}\widetilde{\mathbf{X}}^\top\widetilde{\mathbf{W}}\widetilde{\boldsymbol{\xi}}\| > l_n) \leq \mathbb{P}([\sqrt{\frac{(q+1)p}{n}}R^{3/2}]^4(\sup_{\|\boldsymbol{u}\|=1}|\boldsymbol{u}^\top\widetilde{\boldsymbol{\xi}}|^2)^2 > l_n^4)$$

$$\leq \frac{(q+1)^2p^2R^6\mathbb{E}(\sup_{\|\boldsymbol{u}\|=1}|\boldsymbol{u}^\top\widetilde{\boldsymbol{\xi}}|^2)^2}{n^2l_n^4}$$

$$\leq KR^6\frac{(q+1)^4p^4}{n^2l_n^4} \to 0,$$

where the last inequality follows the same proof of Lemma 3 in Sofer et al. (2017) by setting $\delta = 2$. This completes the proof. □

#### 2. Additional simulation results.

2.1. *Consistency under the null.* To evaluate the performance of our proposed estimators under the null hypothesis of no mediation effect, we conducted additional simulation studies. These settings follow the same framework as described in Section 4.1 of the main text, with one key modification: the cumulative variance component for the true mediators was set to $\sigma_{11}^2 = 0$, thereby eliminating any indirect effect. All $p = 2000$ mediators were non-mediators, and we considered three scenarios that varied the composition of non-mediators while preserving their total number: (S1) 600 $\boldsymbol{M}_{\mathcal{I}_1}$, 600 $\boldsymbol{M}_{\mathcal{I}_2}$, and 800 $\boldsymbol{M}_{\mathcal{I}_3}$; (S2) 1200 $\boldsymbol{M}_{\mathcal{I}_1}$, 600 $\boldsymbol{M}_{\mathcal{I}_2}$, and 200 $\boldsymbol{M}_{\mathcal{I}_3}$; and (S3) 600 $\boldsymbol{M}_{\mathcal{I}_1}$, 1200 $\boldsymbol{M}_{\mathcal{I}_2}$, and 200 $\boldsymbol{M}_{\mathcal{I}_3}$. The results are shown in Table S1. Across all settings, both $\hat{R}_{\text{med; causal}}$ and $\hat{Q}^2_{\text{med}}$ consistently approached zero and all performance metrics were on a comparable scale to those reported in Table 1 of the main text.

TABLE S1

*Performance of proposed estimators under the null. Bias represents empirical bias, SD represents standard deviation, MAD is mean absolute deviation, and MSE is mean square error, all based on 100 repetitions.*

| | | $R^2_{\mathrm{med;causal}}$ | | | | $Q^2_{\mathrm{med}}$ | | | |
|---|---|---|---|---|---|---|---|---|---|
| Scenario | $n$ | Bias | SD | MAD | MSE | Bias | SD | MAD | MSE |
| S1 | 500 | 0.0165 | 0.0294 | 0.0276 | 0.0009 | 0.029 | 0.195 | 0.152 | 0.0377 |
| | 1000 | 0.0033 | 0.014 | 0.0115 | 0.0002 | 0.0078 | 0.078 | 0.0622 | 0.006 |
| | 1500 | 0.0018 | 0.0085 | 0.0066 | 0.0001 | 0.0054 | 0.0499 | 0.0365 | 0.0025 |
| | 2000 | 0.0031 | 0.0069 | 0.0061 | 0 | 0.0135 | 0.0348 | 0.0303 | 0.0012 |
| S2 | 500 | 0.0166 | 0.0408 | 0.0339 | 0.0016 | 0.0655 | 0.255 | 0.18 | 0.0642 |
| | 1000 | 0.0102 | 0.0198 | 0.0175 | 0.0004 | 0.0293 | 0.107 | 0.0871 | 0.0114 |
| | 1500 | 0.0047 | 0.0125 | 0.0106 | 0.0002 | 0.0175 | 0.0631 | 0.053 | 0.0039 |
| | 2000 | 0.0033 | 0.0105 | 0.0083 | 0.0001 | 0.0119 | 0.0553 | 0.0414 | 0.003 |
| S3 | 500 | 0.0077 | 0.0264 | 0.0216 | 0.0007 | -0.149 | 1.16 | 0.274 | 1.34 |
| | 1000 | 0.0067 | 0.0146 | 0.0125 | 0.0002 | 0.0235 | 0.077 | 0.0618 | 0.0059 |
| | 1500 | 0.0015 | 0.0087 | 0.007 | 0.0001 | 0.0043 | 0.0436 | 0.0337 | 0.0019 |
| | 2000 | 0.0032 | 0.0064 | 0.0058 | 0 | 0.0141 | 0.0311 | 0.0282 | 0.001 |

2.2. *Inference under the null.* In addition to point estimation, we assessed the inferential validity of our jackknife-based confidence interval procedure under the null hypothesis. Using scenario S1 with $n = 500$ and $p = 2000$ as a representative setting, we estimated the empirical coverage rate of the 95% confidence interval for the total mediation effect. Across 1000 replicates, the interval captured the true null value (zero) 992 times, yielding a coverage probability of 99.2%. This result indicates that the proposed confidence interval is conservative yet effective in maintaining nominal coverage levels under the null.

2.3. *Computation speed and performance under varying population prevalence.* We presented the computational time of our proposed method compared to existing approaches and examined performance under varying population prevalence settings. Table S2 presents computation times for different methods across various sample sizes and population prevalence values. Tables S3 and S4 show the relative performance of different estimators under varying population prevalence settings.

TABLE S2

*Computation speed for different approaches (in seconds). Each cell presents the average running time, with the standard deviation of the running time provided in parentheses.*

| | Proposed | | HIMA | | HDMA | | BAMA | |
|---|---|---|---|---|---|---|---|---|
| $n$, $K$ | 0.05 | 0.5 | 0.05 | 0.5 | 0.05 | 0.5 | 0.05 | 0.5 |
| 500 | 12.92 | 25.74 | 2.44 | 2.64 | 4.78 | 5.02 | 19.03 | 18.92 |
| | (0.21) | (0.74) | (0.12) | (0.35) | (0.1) | (0.56) | (2.72) | (3.15) |
| 1000 | 15.79 | 16.8 | 3.46 | 4.58 | 9.67 | 12.28 | 37.44 | 39.29 |
| | (0.25) | (0.67) | (0.17) | (1.48) | (0.23) | (4.2) | (6.62) | (7.35) |
| 1500 | 19.62 | 21 | 4.75 | 5.23 | 17.73 | 18.79 | 57.06 | 58.56 |
| | (0.28) | (0.75) | (0.31) | (0.63) | (0.38) | (1.26) | (10.25) | (10.54) |
| 2000 | 23.85 | 25.59 | 6.33 | 8.19 | 30.87 | 36.82 | 78.71 | 81.34 |
| | (0.27) | (0.72) | (0.49) | (2.55) | (0.71) | (8.77) | (13.65) | (14.07) |

TABLE S3

*Comparison of relative performance of IE estimators. Each cell reports the relative bias, with the standard deviation of the relative deviation shown in parentheses.*

| | | $R^2_{\text{med;causal}}$ Measure | | | | Method-Specific Measure | | |
|---|---|---|---|---|---|---|---|---|
| K | n | Proposed | HIMA | HDMA | BAMA | HIMA | HDMA | BAMA |
| 0.2 | 500 | 0.13 | -0.2 | 0.62 | -0.9 | 0.63 | 0.92 | -0.92 |
| | | (0.39) | (1.32) | (1.45) | (0.01) | (0.58) | (2.06) | (0.01) |
| | 1000 | 0.02 | 0.14 | 1.12 | -0.92 | 0.78 | 2.56 | -0.93 |
| | | (0.22) | (0.98) | (0.79) | (0.01) | (0.41) | (2.18) | (0.01) |
| | 1500 | 0.02 | 0.07 | 1.01 | -0.92 | 0.78 | 2.17 | -0.93 |
| | | (0.15) | (0.77) | (0.75) | (0.01) | (0.34) | (1.89) | (0.01) |
| | 2000 | 0.02 | -0.05 | 1.10 | -0.92 | 0.78 | 2.48 | -0.93 |
| | | (0.12) | (0.66) | (0.59) | (0.01) | (0.28) | (1.74) | (0.01) |
| 0.5 | 500 | 0.06 | -0.22 | 0.62 | -0.9 | 0.42 | 0.98 | -0.92 |
| | | (0.49) | (1.27) | (1.44) | (0.01) | (0.51) | (1.89) | (0.01) |
| | 1000 | 0.05 | -0.06 | 0.87 | -0.92 | 0.55 | 1.96 | -0.93 |
| | | (0.23) | (0.95) | (0.85) | (0.01) | (0.38) | (1.91) | (0.01) |
| | 1500 | 0.06 | -0.08 | 0.94 | -0.92 | 0.51 | 1.89 | -0.93 |
| | | (0.17) | (0.79) | (0.69) | (0.01) | (0.31) | (1.73) | (0.01) |
| | 2000 | 0.04 | -0.12 | 0.79 | -0.93 | 0.58 | 1.71 | -0.93 |
| | | (0.13) | (0.58) | (0.59) | (0.01) | (0.29) | (1.97) | (0.01) |

TABLE S4
*Comparison of relative performance of proportion-mediated estimators. Each cell reports the relative bias, with the standard deviation of the relative deviation shown in parentheses.*

| | | $Q^2_{med}$ Measure | | | | Method-Specific Measure | |
|---|---|---|---|---|---|---|---|
| K | n | Proposed | HIMA | HDMA | BAMA | HIMA | HDMA |
| 0.2 | 500 | 0.13 | -0.59 | -0.03 | -0.12 | 0.01 | 0.2 |
| | | (0.35) | (0.65) | (0.92) | (0.2) | (0.37) | (1.3) |
| | 1000 | 0.07 | -0.33 | 0.65 | -0.24 | 0.10 | 1.18 |
| | | (0.23) | (0.54) | (0.73) | (0.14) | (0.26) | (1.31) |
| | 1500 | 0.05 | -0.35 | 0.58 | -0.27 | 0.08 | 0.93 |
| | | (0.18) | (0.46) | (0.74) | (0.11) | (0.22) | (1.17) |
| | 2000 | 0.06 | -0.40 | 0.75 | -0.28 | 0.10 | 1.15 |
| | | (0.14) | (0.41) | (0.71) | (0.09) | (0.19) | (1.09) |
| 0.5 | 500 | 0.02 | -0.57 | 0.07 | -0.14 | -0.06 | 0.33 |
| | | (0.42) | (0.65) | (0.97) | (0.2) | (0.35) | (1.3) |
| | 1000 | 0.07 | -0.41 | 0.50 | -0.23 | 0.03 | 0.95 |
| | | (0.25) | (0.57) | (0.78) | (0.16) | (0.28) | (1.28) |
| | 1500 | 0.08 | -0.41 | 0.62 | -0.28 | 0.01 | 0.93 |
| | | (0.16) | (0.48) | (0.74) | (0.1) | (0.21) | (1.18) |
| | 2000 | 0.08 | -0.40 | 0.55 | -0.29 | 0.05 | 0.80 |
| | | (0.15) | (0.38) | (0.74) | (0.1) | (0.21) | (1.3) |

2.4. *Robustness under strong mediation signals.* To assess robustness of our method under stronger mediation signals, we conducted an additional simulation study that extended the setting used in the main text (Section 5.1). The goal of this simulation is to examine how the proposed method performs when there is a varying number of true mediators with stronger effects. Specifically, all settings remained the same as in Section 5.1, except that we varied the number of true mediators to simulate stronger signal scenarios by fixing $\sigma^2_\beta$.

In this setting, we fixed the proportion of non-zero $\alpha$ effects at 20%, corresponding to 400 out of 2000 candidate mediators having non-zero exposure–mediator effects. We then varied the number of mediators that had non-zero $\beta$ effects (mediator–outcome effects) such that the true number of mediators (i.e., those with both $\alpha \neq 0$ and $\beta \neq 0$) was 50, 20, 10, and 5, respectively. This implies increasingly stronger signals per mediator as the number of true mediators decreases while the total mediation effect is fixed.

Table S5 presents the results across varying numbers of true mediators. Overall, our method exhibited satisfactory performance in scenarios with 10 or more true mediators, with small bias, standard deviation, and MAD in estimating both $R^2_{\text{med;causal}}$ and $Q^2_{\text{med}}$. When the number of true mediators was as low as 5, the bias became slightly more pronounced relative to the true values, reflecting some finite-sample limitations for our estimation procedure under extremely sparse and strong signal settings.

TABLE S5

*Performance of proposed estimators across varying number of true mediators with strong effect. True values for $R^2_{\text{med;causal}}$ and $Q^2_{\text{med}}$ are fixed at 0.1071 and 0.375 respectively.*

| # of Mediators | $n$ | $R^2_{\text{med;causal}}$ | | | | $Q^2_{\text{med}}$ | | | |
|---|---|---|---|---|---|---|---|---|---|
| | | Bias | SD | MAD | MSE | Bias | SD | MAD | MSE |
| 5 | 500 | -0.0156 | 0.0539 | 0.047 | 0.0031 | -0.0616 | 0.179 | 0.162 | 0.0356 |
| | 1000 | -0.0205 | 0.0463 | 0.0422 | 0.0025 | -0.0702 | 0.169 | 0.155 | 0.0333 |
| | 1500 | -0.0173 | 0.0408 | 0.0373 | 0.0019 | -0.0546 | 0.148 | 0.133 | 0.0247 |
| | 2000 | -0.0153 | 0.0414 | 0.0363 | 0.0019 | -0.0458 | 0.155 | 0.132 | 0.0259 |
| 10 | 500 | -0.001 | 0.0494 | 0.0394 | 0.0024 | -0.018 | 0.164 | 0.135 | 0.0268 |
| | 1000 | -0.009 | 0.035 | 0.03 | 0.0013 | -0.0212 | 0.139 | 0.115 | 0.0197 |
| | 1500 | -0.0113 | 0.0308 | 0.0272 | 0.0011 | -0.0333 | 0.126 | 0.108 | 0.0168 |
| | 2000 | -0.008 | 0.0297 | 0.0251 | 0.0009 | -0.0188 | 0.114 | 0.0933 | 0.0133 |
| 20 | 500 | -0.0066 | 0.0408 | 0.0324 | 0.0017 | -0.0215 | 0.153 | 0.12 | 0.0237 |
| | 1000 | -0.0103 | 0.0263 | 0.0232 | 0.0008 | -0.0282 | 0.101 | 0.0889 | 0.0109 |
| | 1500 | -0.0095 | 0.0232 | 0.0201 | 0.0006 | -0.0408 | 0.0853 | 0.0739 | 0.0089 |
| | 2000 | -0.0122 | 0.0225 | 0.0203 | 0.0007 | -0.0415 | 0.0874 | 0.0799 | 0.0093 |
| 50 | 500 | -0.0001 | 0.0351 | 0.0282 | 0.0012 | -0.0068 | 0.136 | 0.109 | 0.0184 |
| | 1000 | -0.0079 | 0.0242 | 0.021 | 0.0006 | -0.0188 | 0.104 | 0.0863 | 0.0111 |
| | 1500 | -0.0079 | 0.0204 | 0.0175 | 0.0005 | -0.0205 | 0.0767 | 0.0638 | 0.0062 |
| | 2000 | -0.0089 | 0.0188 | 0.0167 | 0.0004 | -0.0223 | 0.0752 | 0.063 | 0.0061 |

2.5. *Robustness to misspecified disease prevalence.* To evaluate the sensitivity of our estimator to the misspecification of the population disease prevalence used in the estimation procedure, we conducted an additional simulation study in which the true prevalence was fixed at $K = 0.05$, while the assumed prevalence used in estimation, denoted by $K_{\text{est}}$, was varied across a plausible range $\{0.01, 0.02, \ldots, 0.09\}$. This setup reflects practical situations in case-control studies where the disease prevalence is estimated from a large sample that reflects the study population rather than known. All other simulation settings were kept identical to scenario 1 used in the simulation study reported in Section 5.1 of the main text.

Table S6 summarizes the empirical performance of the proposed estimators across different sample sizes and assumed prevalence values. Overall, the proposed method demonstrated reasonable robustness to moderate misspecification of $K$. In particular, when $K_{\text{est}}$ was close to the true value (e.g., within the range 0.04–0.06), the bias and MSE of both $R^2_{\text{med;causal}}$ and $Q^2_{\text{med}}$ remained small across all sample sizes considered. As the deviation between $K_{\text{est}}$ and the true prevalence increased, the bias became more noticeable, especially for smaller sample sizes. Nevertheless, the overall estimation accuracy remained stable and improved with increasing sample size. These results suggest that while accurate specification of disease prevalence is desirable, the proposed procedure is relatively tolerant to small-to-moderate misspecification in practice.

TABLE S6
*Performance of proposed estimators under varying $K_{est}$ values with true $K = 0.05$. True values for $R^2_{\text{med;causal}}$ and $Q^2_{\text{med}}$ are 0.1071 and 0.375 respectively.*

| $n$ | $K_{\text{est}}$ | $R^2_{\text{med;causal}}$ | | | | $Q^2_{\text{med}}$ | | | |
|---|---|---|---|---|---|---|---|---|---|
| | | Bias | SD | MAD | MSE | Bias | SD | MAD | MSE |
| | 0.01 | -0.0307 | 0.0253 | 0.0337 | 0.0016 | -0.0353 | 0.111 | 0.091 | 0.0134 |
| | 0.02 | -0.0197 | 0.0285 | 0.0277 | 0.0012 | -0.0311 | 0.11 | 0.0892 | 0.013 |
| | 0.03 | -0.012 | 0.0303 | 0.0255 | 0.0011 | -0.0279 | 0.11 | 0.0874 | 0.0128 |
| | 0.04 | -0.0057 | 0.0322 | 0.0254 | 0.0011 | -0.0237 | 0.111 | 0.0868 | 0.0128 |
| 500 | 0.05 | -0.0004 | 0.0335 | 0.0259 | 0.0011 | -0.0211 | 0.112 | 0.0858 | 0.0128 |
| | 0.06 | 0.0044 | 0.035 | 0.0276 | 0.0012 | -0.0174 | 0.112 | 0.0859 | 0.0127 |
| | 0.07 | 0.0084 | 0.0355 | 0.0286 | 0.0013 | -0.0148 | 0.11 | 0.0838 | 0.0123 |
| | 0.08 | 0.0124 | 0.0365 | 0.0303 | 0.0015 | -0.0117 | 0.11 | 0.0838 | 0.0122 |
| | 0.09 | 0.0154 | 0.0376 | 0.032 | 0.0016 | -0.0102 | 0.112 | 0.0849 | 0.0126 |
| | 0.01 | -0.0322 | 0.0129 | 0.0322 | 0.0012 | -0.0036 | 0.102 | 0.0761 | 0.0104 |
| | 0.02 | -0.0214 | 0.0147 | 0.0224 | 0.0007 | -0.0028 | 0.0998 | 0.0738 | 0.0099 |
| | 0.03 | -0.0136 | 0.0159 | 0.0169 | 0.0004 | -0.0006 | 0.0984 | 0.0723 | 0.0096 |
| | 0.04 | -0.0075 | 0.0168 | 0.0145 | 0.0003 | 0.0013 | 0.096 | 0.0706 | 0.0091 |
| 1000 | 0.05 | -0.0026 | 0.0177 | 0.0141 | 0.0003 | 0.0028 | 0.0954 | 0.0704 | 0.009 |
| | 0.06 | 0.0021 | 0.0183 | 0.0146 | 0.0003 | 0.0053 | 0.0945 | 0.0699 | 0.0089 |
| | 0.07 | 0.0065 | 0.0189 | 0.0159 | 0.0004 | 0.0082 | 0.0932 | 0.0689 | 0.0087 |
| | 0.08 | 0.0103 | 0.0198 | 0.0172 | 0.0005 | 0.0105 | 0.0928 | 0.0682 | 0.0086 |
| | 0.09 | 0.0137 | 0.0199 | 0.0187 | 0.0006 | 0.0124 | 0.0915 | 0.0675 | 0.0084 |
| | 0.01 | -0.0367 | 0.0128 | 0.0367 | 0.0015 | -0.0342 | 0.0633 | 0.0574 | 0.0051 |
| | 0.02 | -0.0265 | 0.0143 | 0.0269 | 0.0009 | -0.0329 | 0.0617 | 0.0558 | 0.0049 |
| | 0.03 | -0.0193 | 0.0153 | 0.0206 | 0.0006 | -0.0305 | 0.0609 | 0.0542 | 0.0046 |
| | 0.04 | -0.0135 | 0.016 | 0.017 | 0.0004 | -0.0282 | 0.0603 | 0.0527 | 0.0044 |
| 1500 | 0.05 | -0.0087 | 0.0167 | 0.0151 | 0.0004 | -0.0259 | 0.0602 | 0.0516 | 0.0043 |
| | 0.06 | -0.0044 | 0.0171 | 0.0141 | 0.0003 | -0.0234 | 0.0595 | 0.0503 | 0.0041 |
| | 0.07 | -0.0005 | 0.0177 | 0.0144 | 0.0003 | -0.0208 | 0.0597 | 0.0499 | 0.004 |
| | 0.08 | 0.003 | 0.0182 | 0.015 | 0.0003 | -0.0184 | 0.0597 | 0.0495 | 0.0039 |
| | 0.09 | 0.006 | 0.0184 | 0.0157 | 0.0004 | -0.0166 | 0.0592 | 0.0491 | 0.0037 |
| | 0.01 | -0.0367 | 0.0103 | 0.0368 | 0.0015 | -0.0258 | 0.061 | 0.0519 | 0.0044 |
| | 0.02 | -0.0266 | 0.0119 | 0.027 | 0.0008 | -0.026 | 0.0594 | 0.051 | 0.0042 |
| | 0.03 | -0.0194 | 0.0128 | 0.0202 | 0.0005 | -0.0244 | 0.0587 | 0.0502 | 0.004 |
| | 0.04 | -0.0137 | 0.0135 | 0.0156 | 0.0004 | -0.0225 | 0.0578 | 0.049 | 0.0038 |
| 2000 | 0.05 | -0.0089 | 0.0139 | 0.0129 | 0.0003 | -0.0206 | 0.057 | 0.0482 | 0.0036 |
| | 0.06 | -0.0047 | 0.0146 | 0.0121 | 0.0002 | -0.0186 | 0.0571 | 0.0477 | 0.0036 |
| | 0.07 | -0.0007 | 0.015 | 0.0119 | 0.0002 | -0.0161 | 0.057 | 0.0468 | 0.0035 |
| | 0.08 | 0.0027 | 0.0155 | 0.0123 | 0.0002 | -0.0143 | 0.0569 | 0.0461 | 0.0034 |
| | 0.09 | 0.0059 | 0.016 | 0.0132 | 0.0003 | -0.0122 | 0.0568 | 0.0456 | 0.0033 |

2.6. *Sensitivity of $Q^2_{med}$ to small total effects.* To investigate the stability of the relative mediation effect measure $Q^2_{\text{med}}$ under small total effects, we conducted a simulation study in which the total effect of the exposure on the outcome, defined as $\sigma^2_\alpha \sigma^2_{11} + \gamma^2$, was varied from a moderate magnitude to values close to zero. All other simulation settings were identical to those used in scenario 1 of Section 5.1 in the main text, with targeted modifications to reduce the total effect size. Specifically, we varied the variance components $\sigma^2_\alpha$, $\sigma^2_{11}$, and the direct effect parameter $\gamma$ proportionally by a scaling factor. This controlled reduction in total effect while keeping the relative mediation effect $Q^2_{\text{med}}$ fixed at 0.8333 across all scenarios. This setup reflects scenarios where mediation explains a large proportion of a weak total effect, a condition under which the ratio-based $Q^2_{\text{med}}$ may become unstable and difficult to interpret.

Table S7 presents the simulation results. As expected, when the total effect was very small (e.g., $< 0.003$), we observed substantial bias and variability in the estimated $Q^2_{\text{med}}$, particularly for smaller sample sizes. This instability diminished gradually as the total effect increased. These findings indicate that $Q^2_{\text{med}}$ should be interpreted with caution when the total effect is weak, even if the true relative total mediation effect appears relatively strong. While the performance of $R^2_{\text{med;causal}}$ was notably better in comparison, it also exhibited a non-negligible bias when both the total effect and sample size were small (e.g., $n = 500$). However, this bias decreased substantially as the sample size increased, and the estimator became more stable and accurate. These findings indicated that the observed instability was primarily driven by the ratio-based structure of $Q^2_{\text{med}}$, while $R^2_{\text{med;causal}}$ remained more robust, especially in larger samples.

TABLE S7
*Performance of proposed estimators under varying small total mediation effect levels. The true values for $Q^2_{\text{med}}$ is fixed at 0.8333.*

| $n$ | Total effect | $R^2_{\text{med;causal}}$ | | | | $Q^2_{\text{med}}$ | | | |
|---|---|---|---|---|---|---|---|---|---|
| | | Bias | SD | MAD | MSE | Bias | SD | MAD | MSE |
| 500 | 0.00096 | 0.0025 | 0.0034 | 0.0029 | $< 10^{-4}$ | -0.553 | 0.2636 | 0.558 | 0.3746 |
| | 0.00192 | 0.003 | 0.0042 | 0.0036 | $< 10^{-4}$ | -0.4648 | 0.2531 | 0.4715 | 0.2795 |
| | 0.00287 | 0.0034 | 0.004 | 0.004 | $< 10^{-4}$ | -0.356 | 0.2499 | 0.3765 | 0.1886 |
| | 0.00478 | 0.0027 | 0.0048 | 0.0042 | $< 10^{-4}$ | -0.3743 | 0.2525 | 0.3873 | 0.2032 |
| | 0.00951 | 0.003 | 0.0065 | 0.0056 | 0.0001 | -0.2542 | 0.2573 | 0.2856 | 0.1302 |
| | 0.01884 | 0.0034 | 0.0089 | 0.0076 | 0.0001 | -0.1707 | 0.2252 | 0.218 | 0.0793 |
| 1000 | 0.00096 | 0.0004 | 0.001 | 0.0008 | $< 10^{-4}$ | -0.4934 | 0.2718 | 0.5047 | 0.3166 |
| | 0.00192 | 0.0006 | 0.0015 | 0.0012 | $< 10^{-4}$ | -0.3917 | 0.256 | 0.4037 | 0.2183 |
| | 0.00287 | 0.0005 | 0.0017 | 0.0014 | $< 10^{-4}$ | -0.3542 | 0.2618 | 0.3777 | 0.1933 |
| | 0.00478 | 0.0004 | 0.0022 | 0.0018 | $< 10^{-4}$ | -0.2869 | 0.2296 | 0.3052 | 0.1345 |
| | 0.00951 | -0.0001 | 0.0032 | 0.0026 | $< 10^{-4}$ | -0.1983 | 0.2096 | 0.2324 | 0.0828 |
| | 0.01884 | -0.0006 | 0.0041 | 0.0032 | $< 10^{-4}$ | -0.0938 | 0.1506 | 0.1386 | 0.0313 |
| 1500 | 0.00096 | 0.0002 | 0.0007 | 0.0006 | $< 10^{-4}$ | -0.434 | 0.2372 | 0.4366 | 0.244 |
| | 0.00192 | 0.0003 | 0.001 | 0.0008 | $< 10^{-4}$ | -0.2795 | 0.2209 | 0.2961 | 0.1264 |
| | 0.00287 | 0.0001 | 0.001 | 0.0008 | $< 10^{-4}$ | -0.2497 | 0.2295 | 0.2763 | 0.1145 |
| | 0.00478 | $< 10^{-4}$ | 0.0014 | 0.0012 | $< 10^{-4}$ | -0.1887 | 0.2003 | 0.2193 | 0.0753 |
| | 0.00951 | -0.0001 | 0.0021 | 0.0017 | $< 10^{-4}$ | -0.0911 | 0.1679 | 0.1497 | 0.0362 |
| | 0.01884 | -0.0011 | 0.0034 | 0.0031 | $< 10^{-4}$ | -0.073 | 0.1363 | 0.1218 | 0.0237 |
| 2000 | 0.00096 | 0.0001 | 0.0005 | 0.0004 | $< 10^{-4}$ | -0.3523 | 0.2351 | 0.3625 | 0.1789 |
| | 0.00192 | 0.0001 | 0.0007 | 0.0005 | $< 10^{-4}$ | -0.2298 | 0.213 | 0.2545 | 0.0977 |
| | 0.00287 | 0.0001 | 0.001 | 0.0008 | $< 10^{-4}$ | -0.2224 | 0.1995 | 0.2418 | 0.0889 |
| | 0.00478 | $< 10^{-4}$ | 0.0011 | 0.0009 | $< 10^{-4}$ | -0.1072 | 0.1556 | 0.145 | 0.0355 |
| | 0.00951 | -0.0001 | 0.0017 | 0.0014 | $< 10^{-4}$ | -0.0872 | 0.1384 | 0.128 | 0.0266 |
| | 0.01884 | -0.001 | 0.0026 | 0.0022 | $< 10^{-4}$ | -0.0547 | 0.1252 | 0.1077 | 0.0185 |

2.7. *Sensitivity to FDR threshold in screening step.* To assess the sensitivity of our method to the choice of false discovery rate (FDR) threshold in the variable screening step, we conducted a simulation study where the FDR cutoff was varied from 0.01 to 0.25. This range encompasses both conservative and more liberal screening settings. All other simulation parameters were kept the same as those used for the main results reported in Section 5.1, scenario 1.

Table S8 summarizes the estimation performance under different FDR thresholds across multiple sample sizes. We found that the proposed estimator was relatively robust to modest variations in the FDR threshold. In particular, FDR values between 0.01 and 0.05 yielded small bias and stable estimation for both $R^2_{\text{med;causal}}$ and $Q^2_{\text{med}}$. As the FDR cutoff increased to 0.25, bias became more noticeable and estimation variability increased, especially for smaller sample sizes. This is likely due to more noise variables being retained in the screening step, which can introduce spurious signal and inflate estimation error. Based on these results, we recommend using an FDR threshold in the range of 0.01 to 0.05 for practical applications. This range offers a favorable balance between excluding non-mediators and retaining true signals, thereby promoting both estimation accuracy and robustness.

TABLE S8

*Performance of proposed estimators under varying FDR cutoff thresholds across different sample sizes. True values for $R^2_{\text{med;causal}}$ and $Q^2_{\text{med}}$ are 0.1071 and 0.375 respectively.*

| $n$ | FDR cutoff | $R^2_{\text{med;causal}}$ | | | | $Q^2_{\text{med}}$ | | | |
|---|---|---|---|---|---|---|---|---|---|
| | | Bias | SD | MAD | MSE | Bias | SD | MAD | MSE |
| 500 | 0.01 | -0.0004 | 0.0335 | 0.0259 | 0.0011 | -0.0211 | 0.112 | 0.0858 | 0.0128 |
| | 0.05 | -0.0068 | 0.0306 | 0.0248 | 0.001 | -0.0302 | 0.103 | 0.0842 | 0.0114 |
| | 0.1 | -0.0097 | 0.0318 | 0.0252 | 0.0011 | -0.0377 | 0.105 | 0.0846 | 0.0124 |
| | 0.25 | -0.0208 | 0.027 | 0.0277 | 0.0012 | -0.0625 | 0.0934 | 0.089 | 0.0125 |
| 1000 | 0.01 | -0.0026 | 0.0177 | 0.0141 | 0.0003 | 0.0028 | 0.0954 | 0.0704 | 0.009 |
| | 0.05 | -0.0087 | 0.017 | 0.0153 | 0.0004 | -0.012 | 0.0951 | 0.0701 | 0.0091 |
| | 0.1 | -0.0137 | 0.0161 | 0.0176 | 0.0004 | -0.025 | 0.0933 | 0.0736 | 0.0092 |
| | 0.25 | -0.0234 | 0.0154 | 0.0243 | 0.0008 | -0.0503 | 0.0937 | 0.0852 | 0.0112 |
| 1500 | 0.01 | -0.0087 | 0.0167 | 0.0151 | 0.0004 | -0.0259 | 0.0602 | 0.0516 | 0.0043 |
| | 0.05 | -0.0145 | 0.0154 | 0.0173 | 0.0004 | -0.0403 | 0.0584 | 0.0567 | 0.005 |
| | 0.1 | -0.0203 | 0.0148 | 0.0213 | 0.0006 | -0.0557 | 0.0558 | 0.065 | 0.0062 |
| | 0.25 | -0.0297 | 0.0128 | 0.0297 | 0.001 | -0.0812 | 0.0519 | 0.083 | 0.0093 |
| 2000 | 0.01 | -0.0089 | 0.0139 | 0.0129 | 0.0003 | -0.0206 | 0.057 | 0.0482 | 0.0036 |
| | 0.05 | -0.0148 | 0.0134 | 0.0167 | 0.0004 | -0.0357 | 0.0568 | 0.0541 | 0.0045 |
| | 0.1 | -0.0198 | 0.0126 | 0.0204 | 0.0005 | -0.0486 | 0.0562 | 0.0617 | 0.0055 |
| | 0.25 | -0.03 | 0.0112 | 0.0301 | 0.001 | -0.0773 | 0.0529 | 0.0825 | 0.0087 |

2.8. *Robustness to residual correlation among mediators.* To assess the robustness of the proposed method under violations of the assumption of weak residual correlation among mediators, we conducted a simulation study in which residuals from the mediator model were allowed to have dense and non-negligible correlation. Specifically, we constructed the correlation matrix via a latent factor model with three latent factors, where factor loadings were drawn from a zero-mean normal distribution and independent uniform noise was added to the diagonal to ensure positive definiteness. The overall strength of residual correlation was controlled through the standard deviation of the factor loadings and was quantified by the empirical standard deviation of the off-diagonal elements of the resulting correlation matrix (off-diag SD). All other simulation parameters matched those used in the main results presented in Section 5.1 scenario 1.

Table S9 summarizes the estimation performance across varying levels of residual correlation and sample size. We found that the proposed estimator maintained acceptable performance under low to moderate residual correlation (e.g., up to 0.1714 off-diag SD). As off-diag SD increased beyond 0.2481, bias and mean squared error in both $R^2_{\text{med;causal}}$ and $Q^2_{\text{med}}$ became more pronounced, especially for smaller sample sizes. Notably, increasing the sample size mitigated performance degradation, indicating that the estimator remains numerically stable even when the weak correlation assumption is modestly violated.

These results suggest that the proposed approach is reasonably robust to mild residual correlations but may be sensitive under high residual dependence. In practice, we recommend inspecting the empirical residual correlation matrix from the mediator model (after adjusting for exposure and covariates) to assess potential violations. For example, in the WHI data, we observed that the off-diag SD was around 0.17, and the distribution of residual correlations was centered around zero, with approximately 90% of residual correlations falling below 0.25 in absolute value, and only a small proportion exceeding 0.3. Users may calculate the off-diag SD and examine the residual correlation distribution to check whether it is centered near zero with few large values, and consider pre-screening or increasing the number of principal components included in the adjustment step if strong dependencies are observed.

TABLE S9

*Performance of proposed estimators under varying levels of residual correlation strength in mediators across different sample sizes. "Off-diag SD" denotes the empirical standard deviation of off-diagonal elements in the residual correlation matrix after regressing out exposure, covariates, and latent factor $U$.*

| $n$ | Off-diag SD | $R^2_{\text{med;causal}}$ | | | | $Q^2_{\text{med}}$ | | | |
|---|---|---|---|---|---|---|---|---|---|
| | | Bias | SD | MAD | MSE | Bias | SD | MAD | MSE |
| 500 | 0 | 0.0033 | 0.0379 | 0.0304 | 0.0014 | -0.022 | 0.1207 | 0.0979 | 0.0149 |
| | 0.0116 | 0.0036 | 0.0392 | 0.0302 | 0.0015 | -0.0232 | 0.1228 | 0.1009 | 0.0155 |
| | 0.0834 | 0.0135 | 0.043 | 0.0345 | 0.002 | -0.0253 | 0.1176 | 0.0972 | 0.0143 |
| | 0.1714 | 0.0271 | 0.0493 | 0.0443 | 0.0031 | -0.0266 | 0.1227 | 0.0959 | 0.0156 |
| | 0.2481 | 0.0304 | 0.0594 | 0.0526 | 0.0044 | -0.035 | 0.147 | 0.1178 | 0.0226 |
| | 0.3088 | 0.0455 | 0.0654 | 0.0639 | 0.0063 | -0.0362 | 0.1422 | 0.1154 | 0.0213 |
| 1000 | 0 | -0.0049 | 0.0206 | 0.0162 | 0.0004 | -0.0197 | 0.0853 | 0.0666 | 0.0076 |
| | 0.0116 | -0.0036 | 0.0193 | 0.0156 | 0.0004 | -0.0158 | 0.086 | 0.0642 | 0.0076 |
| | 0.0834 | 0.0035 | 0.0234 | 0.0177 | 0.0006 | -0.0195 | 0.0776 | 0.0638 | 0.0063 |
| | 0.1714 | 0.0099 | 0.0315 | 0.0257 | 0.0011 | -0.0342 | 0.0946 | 0.0758 | 0.01 |
| | 0.2481 | 0.0224 | 0.0324 | 0.0313 | 0.0015 | -0.0393 | 0.0818 | 0.0694 | 0.0082 |
| | 0.3088 | 0.0287 | 0.0392 | 0.0374 | 0.0023 | -0.0472 | 0.0958 | 0.0899 | 0.0113 |
| 1500 | 0 | -0.0076 | 0.0173 | 0.0152 | 0.0004 | -0.0297 | 0.0667 | 0.0591 | 0.0053 |
| | 0.0116 | -0.0041 | 0.0151 | 0.0125 | 0.0002 | -0.0109 | 0.0664 | 0.0515 | 0.0045 |
| | 0.0834 | -0.0009 | 0.0204 | 0.0163 | 0.0004 | -0.0332 | 0.0741 | 0.0643 | 0.0065 |
| | 0.1714 | 0.0106 | 0.0206 | 0.0181 | 0.0005 | -0.0319 | 0.0626 | 0.0547 | 0.0049 |
| | 0.2481 | 0.025 | 0.0291 | 0.0308 | 0.0015 | -0.0282 | 0.0831 | 0.0672 | 0.0076 |
| | 0.3088 | 0.0316 | 0.032 | 0.0363 | 0.002 | -0.0399 | 0.0788 | 0.072 | 0.0077 |
| 2000 | 0 | -0.0114 | 0.0159 | 0.016 | 0.0004 | -0.0371 | 0.0632 | 0.0611 | 0.0053 |
| | 0.0116 | -0.0081 | 0.0138 | 0.0122 | 0.0003 | -0.0351 | 0.0551 | 0.0541 | 0.0042 |
| | 0.0834 | 0.0001 | 0.0166 | 0.0129 | 0.0003 | -0.0304 | 0.0645 | 0.0562 | 0.005 |
| | 0.1714 | 0.0136 | 0.0185 | 0.0187 | 0.0005 | -0.0197 | 0.0613 | 0.05 | 0.0041 |
| | 0.2481 | 0.0196 | 0.0248 | 0.0259 | 0.001 | -0.038 | 0.07 | 0.064 | 0.0063 |
| | 0.3088 | 0.0287 | 0.0302 | 0.0339 | 0.0017 | -0.0453 | 0.0788 | 0.0741 | 0.0082 |

2.9. *Confidence interval coverage and computational considerations.* To evaluate the reliability and computational feasibility of confidence interval construction in high-dimensional settings, we conducted additional simulations to evaluate the empirical coverage properties of the proposed jackknife-based inference procedure and to compare it with existing high-dimensional mediation methods.

We first evaluated the empirical coverage of 95% confidence intervals for competing methods, including HIMA, HDMA, and BAMA. The simulation settings followed those used in scenario 1 of Section 5.1 in the main text, with the sample size fixed at $n = 500$ and the number of mediators fixed at $p = 2000$. For HIMA and HDMA, confidence intervals were constructed using bootstrap procedures, while for BAMA, posterior credible intervals were used. The proposed method achieved empirical coverage rates of 96% for both $R^2_{\text{med;causal}}$ and $Q^2_{\text{med}}$. In comparison, HIMA achieved coverage rates of 98% for $R^2_{\text{med;causal}}$ and 86% for $Q^2_{\text{med}}$, while HDMA achieved coverage rates of 96% and 100%, respectively. In contrast, BAMA exhibited zero empirical coverage for both measures under this setting.

Additionally, we assessed the coverage performance of the jackknife confidence intervals for our proposed method under increasingly high-dimensional mediator settings. After increasing the mediator dimension to $p = 5000$, the coverage remained high, at 98% for $R^2_{\text{med;causal}}$ and 94% for $Q^2_{\text{med}}$, indicating that the jackknife procedure maintains satisfactory frequentist properties even in truly high-dimensional settings.

All simulations were conducted on the Minnesota Supercomputing Institute (MSI) interactive partition using a single compute node with one CPU core and 1 GB of memory, without parallelization. The average runtime was approximately 1,207 seconds for $p = 2000$ and 4,161 seconds for $p = 5000$. Although the jackknife procedure is computationally intensive, it is inherently parallelizable because each leave-one-out replicate can be computed independently. In our R package, we optimized the jackknife CI computation by reusing the genetic correlation matrix to avoid redundant matrix multiplications and provided a parallelization option to further reduce runtime. This makes the proposed inference procedure scalable to omics applications involving tens of thousands of mediators.

Finally, we note that traditional product-of-coefficients–based approaches may still be preferable in settings where mediation effects are sparse and strong, and the primary objective is identification of individual mediators. In contrast, the proposed variance-based approach is designed to provide stable and interpretable inference for the total mediation effect, particularly in high-dimensional settings with weak, dense, or bidirectional mediation effects, where product-based methods may suffer from instability or cancellation.

**3. PCGC algorithm.** In the presence of covariates, PCGC regression follows a two-step procedure to estimate $\sigma^2_{11}$. First, the conditional probability of the trait for the $i$-th individual $\hat{P}_i = P(Y_i = 1|X_i, \boldsymbol{C}_i, \boldsymbol{W}_i)$ is estimated using a logistic regression model by regressing $Y$ on $X$, $\boldsymbol{C}$ and $\boldsymbol{W}$. Based on this estimate, we compute the adjusted trait prevalence $\hat{K}_i$ and corresponding liability threshold $\hat{t}_i$ for the $i$-th individual as:

$$\hat{K}_i = \frac{\frac{K(1-P)}{P(1-K)}\hat{P}_i}{1 + \frac{K(1-P)}{P(1-K)}\hat{P}_i - \hat{P}_i},$$

$$\hat{t}_i = \Phi^{-1}(1 - \hat{K}_i),$$

where $\Phi(\cdot)$ denotes the cumulative distribution function (CDF) of the standard normal distribution.

Second, PCGC regression adjusts for ascertainment bias by examining the relationship between phenotypic correlation and correlations among rows of the design matrix, conditioned on the effect of covariates. The phenotypic correlation is expressed as

$$Z_{ij} = \frac{(Y_i - \hat{P}_i)(Y_j - \hat{P}_j)}{\sqrt{\hat{P}_i(1-\hat{P}_i)}\sqrt{\hat{P}_j(1-\hat{P}_j)}},$$

where $Y_i$ and $Y_j$ are the observed binary outcomes for the $i$-th and $j$-th individuals. The correlation matrix, which captures the correlation between individuals based on residuals, is computed as $\hat{G} = \hat{\boldsymbol{\xi}}_{\hat{\mathcal{T}}}\hat{\boldsymbol{\xi}}_{\hat{\mathcal{T}}}^\top/p$. Finally, we obtain an estimator for $\sigma_{11}^2$ by regressing the phenotypic correlation $Z_{ij}$ on the modified correlation:

$$\frac{\varphi(\hat{t}_i)\varphi(\hat{t}_j)\left[1-(\hat{P}_i+\hat{P}_j)\left(\frac{P-K}{P(1-K)}\right)+\hat{P}_i\hat{P}_j\left(\frac{P-K}{P(1-K)}\right)^2\right]\hat{G}_{ij}}{\sqrt{\hat{P}_i(1-\hat{P}_i)}\sqrt{\hat{P}_j(1-\hat{P}_j)}\left(\hat{K}_i+(1-\hat{K}_i)\frac{K(1-P)}{P(1-K)}\right)\left(\hat{K}_j+(1-\hat{K}_j)\frac{K(1-P)}{P(1-K)}\right)}.$$

#### 4. Illustration of cross-fitting procedure.

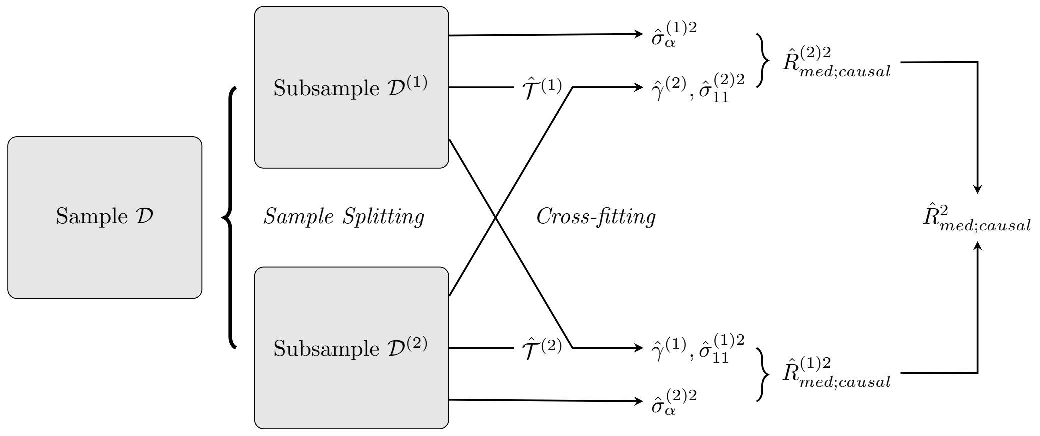


Fig 1: Cross-fitted estimation procedure of $R^2_{\text{med;causal}}$.
